# Sedimentary ancient DNA reveals Late Pleistocene faunal connectivity between Ireland and Eurasia

**DOI:** 10.64898/2026.07.10.737191

**Authors:** Rui Martiniano, Shady Tann-Watson, Taiga McFarlane, Philip Kenny, Allan D. McDevitt, Richard P. Jennings, Helen Lewis, Ruth F. Carden

## Abstract

Late Pleistocene sea-level fluctuations and glacial corridors intermittently connected the island of Ireland to Britain and continental Eurasia, shaping patterns of megafaunal dispersal and occupation. However, the scarcity of genetic data from ancient Irish fauna has limited our understanding of their demographic histories and relationships to continental populations. To address this, we generated sedimentary ancient DNA sequences from Castlepook Cave in southwest Ireland, detecting twelve ancient taxa, including two without zooarchaeological records at the site. We recovered the first mitochondrial sequences from Irish cave hyenas and woolly mammoths, providing new insights into their maternal population history: cave hyenas carried mtDNA haplogroup A1, previously identified in Late Pleistocene European populations, while woolly mammoths belonged to clade III/B2, which was replaced in Europe at approximately the same time. We also identify two mitochondrial clades (1b and 2), consistent with pre-Last Glacial Maximum mitochondrial lineage turnover in Ireland. Together, these results indicate that Irish megafauna formed a biogeographical continuum with their conspecifics in Europe, extend the known geographical range of several mtDNA lineages, and support faunal connectivity at the northwestern edge of Europe prior to the Last Glacial Maximum.

## Introduction

Late Pleistocene Eurasian megafauna experienced pronounced shifts in geographical range, population turnover [1] and extinction [2] in response to climate change and human activity [3]. While these processes are well documented at the continental level [2], their dynamics at the peripheries remain poorly understood.

Despite Ireland’s insular position at the northwestern fringe of Europe, it was intermittently connected to northern Britain and, via southern Britain, to continental Eurasia [4–6], enabling faunal dispersal between landmasses. To date, most genetic research has focused on extant species that colonised Ireland after the Last Glacial Maximum (LGM) period [7,8], either via natural dispersal or introduced by early Mesolithic or Neolithic settlers [9,10]. In contrast, ancient DNA (aDNA) from pre-LGM Irish fauna is very limited, with the notable exception of the brown bear [11]. Consequently, the genetic affinities between Irish Late Pleistocene megafauna and wider Eurasian populations remain unknown. This raises the question of whether Irish megafaunal populations were isolated or formed part of a broader, connected biogeographical network prior to the LGM.

Castlepook Cave (County Cork, Ireland; Fig. 1A) preserves a diverse assemblage of Late Pleistocene vertebrate fauna[12], offering an opportunity to investigate past demography. Early 20th-century excavations recovered approximately 34,000 bones from multiple species, including cave hyena, mammoth, brown bear, reindeer, giant deer, hare, and arctic fox, reflecting Ireland’s Late Pleistocene ecological diversity[13]. Radiocarbon dates place these remains between 54.9-23.3 kya [12], corresponding to the period preceding the Last Glacial Maximum (LGM; c. 26-19 kya), when Ireland is proposed to have been totally or partially covered in ice[14,15].

**Figure 1.**
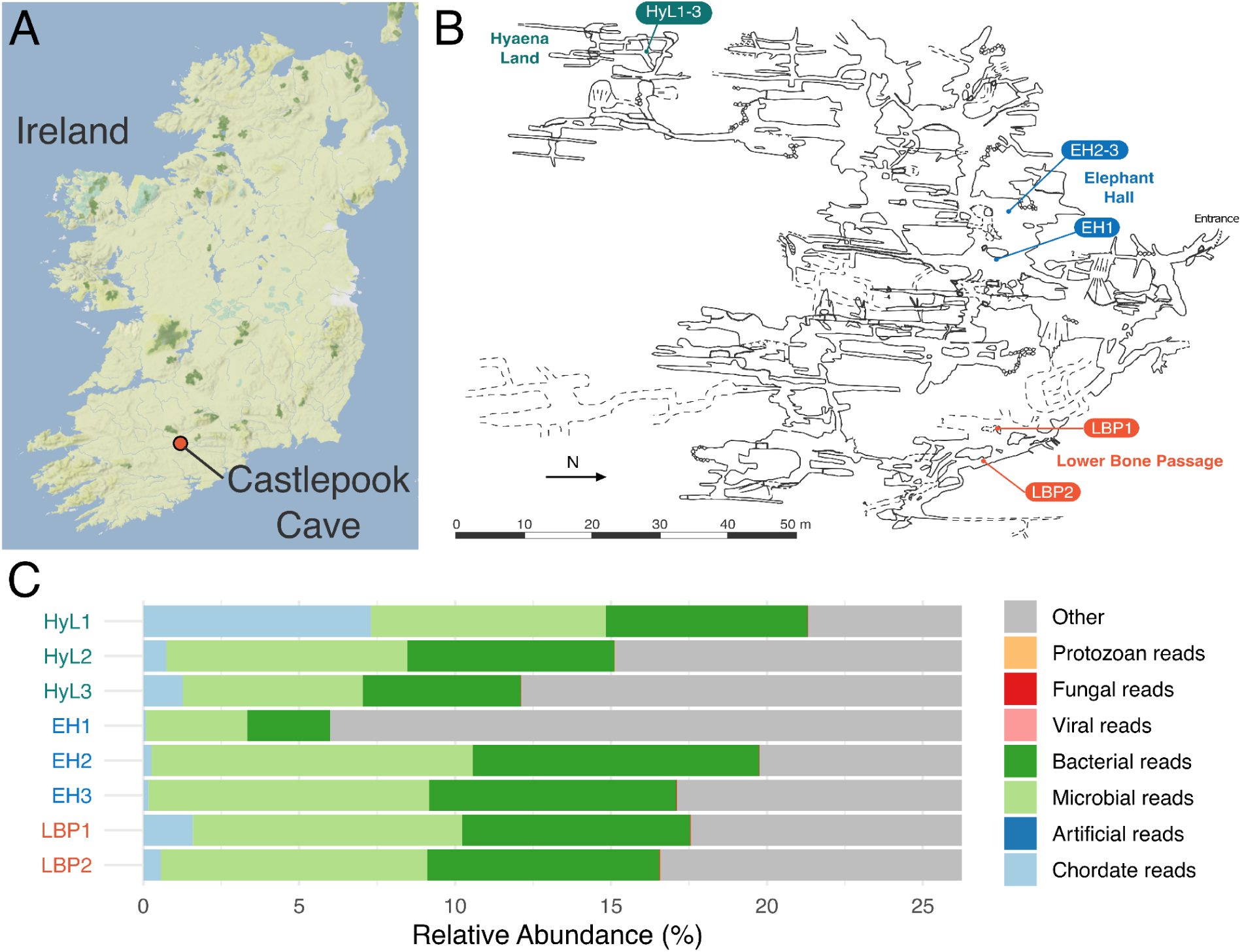
A) Castlepook Cave location. B) Castlepook Cave area and sampling locations. C) High-level taxonomic classification across the different cave areas sampled obtained using kraken. Remaining classified reads not assigned to these major groups were aggregated into the ‘Other’ category comprising low-abundance taxa.

The limited availability of suitable remains from Irish caves for destructive sampling motivated the use of sedimentary ancient DNA (sedaDNA) to recover genetic information directly from the cave sediments. Under favourable conditions, sedaDNA may be preserved for millions of years [16], enabling identification of taxa beyond skeletal morphology and even in the absence of skeletal remains [17], and investigation of genetic affinities [18].

Here, we recovered aDNA from Castlepook Cave sediment samples with the aim of identifying Irish Late Pleistocene fauna and assessing their relationships with Eurasian populations. We detected twelve taxa and reconstructed mitochondrial sequences for some of these ancient species, revealing shared lineages between Irish and continental European fauna prior to the LGM.

## Results

### Overview

We generated sedimentary DNA from eight sediment layers across three areas of Castlepook Cave: three from Hyaena Land (HyL1-3), three from the Elephant Hall (EH1-3) and two from the Lower Bone Passage (LBP1-2) (Fig. 1, Supplementary Table 1). We first assessed metagenomic composition, then identified taxa using aDNA-aware taxonomic classification methods and evaluated their authenticity, and we assessed genetic affinities through mitochondrial phylogenetic placement.

### Sedimentary DNA Recovery and Initial Metagenomic Composition

Following mitochondrial enrichment targeting 30 species (Supplementary Table 2), we generated approximately 1.18 billion paired-end reads across the eight cave deposits sampled. After quality filtering and duplicate removal, ∼11.7 million unique reads (1.21%) were retained, corresponding to ∼1.46 [1.06-2.14] million reads per sample (Supplementary Table 3).

We conducted an initial taxonomic screening with Kraken2[19] to obtain a high-level overview of metagenomic composition across samples. We observed substantial variation between deposits at the level of classified reads, ranging from 3.6% to 15.3% (Supplementary Table 4), with the highest classification rates observed in Hyaena Land (mean = 10.6%). Within the classified reads (Fig. 1C), microbial (3.2-10.3%) and bacterial (2.7-9.2%) sequences dominated the metagenomic profile, whereas chordate DNA was present at low levels across all samples (Supplementary Table 4, mean = 1.75%), with the exception being HyL1 with 7.3%.

Mammalian reads were dominated by a small number of taxa (Supplementary Fig. 4-5), particularly spotted hyena (*Crocuta crocuta*), the closest extant relative, and proxy to the Eurasian cave hyena (*Crocuta crocuta spelaea)[20]*, which is by far the most represented species in our data, followed by *Homo sapiens* and several carnivores and ungulates (Supplementary Table 5 and 6). Taxonomic composition across the cave areas broadly reflected the original excavation finds, with hyena-dominated assemblages in Hyaena Land, and more mixed taxa in Elephant Hall and Lower Bone Passage (Supplementary Table 5) However, this approach also produced spurious assignments (e.g. *Propithecus coquereli*), highlighting the limitations of non-aDNA-aware classification methods for the analysis of degraded sequences[21]. Nevertheless, the results supported the presence of several species represented in the zooarchaeological record of the cave and this encouraged additional investigation.

### Ancient DNA-aware Taxonomic Classification and Authenticity

To improve the accuracy of taxonomic classification, we applied two independent aDNA-aware approaches, quicksand[22] and vegan euka/soibean[23,24], which take into account aDNA-specific characteristics such as post-mortem deamination [25] and short fragment length. Both approaches provided broadly consistent taxonomic profiles, although quicksand was more sensitive, detecting a total of 20 taxa with strong signatures of ancientness and eight with moderate evidence (Figure 2A, Supplementary Table 7-8), whereas euka only detected 17 taxa, of which 15 presented terminal deamination greater than 20% and two taxa with more than 10% Supplementary Figure 6, Supplementary Table 7-8).

**Figure 2.**
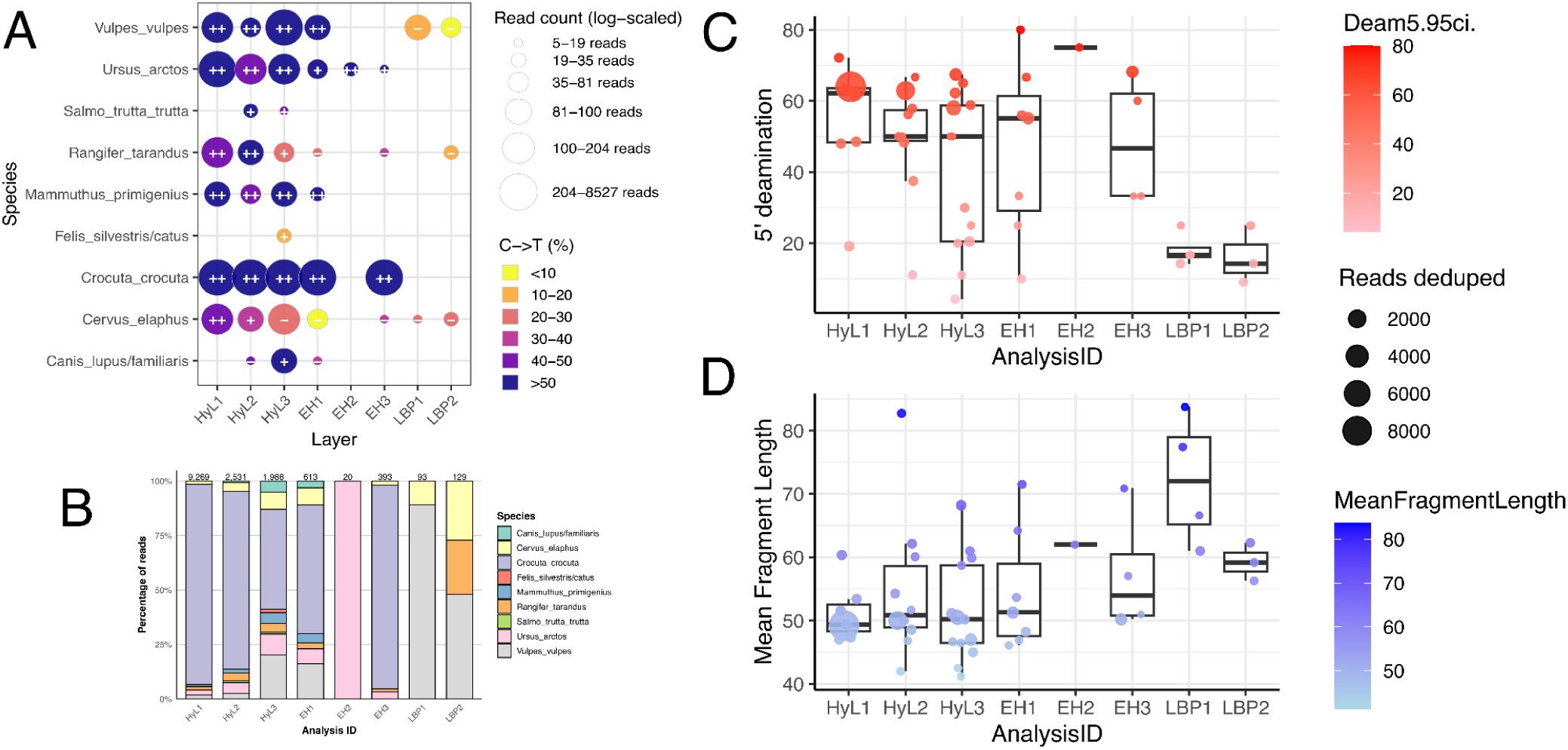
Species detected across different layers and related metrics of authenticity. A) Per species read count, deamination and ancientness classification as determined by quicksand. B) Species composition across all layers, C) Average deamination at the first base of the 5’-end of sequencing reads and D) Average fragment length of sequencing reads in detected taxa in each layer.

The strongest signals in our data belonged to cave hyena (Supplementary Fig. 7) and brown bear (*Ursus arctos;* Supplementary Fig. 8), which were detected widely by both methods in the Hyaena Land and Elephant Hall sediments, with high deamination patterns (>40%), supporting the presence of authentic aDNA (Fig. 2). Additional taxa with authentic aDNA signatures include red fox (*Vulpes vulpes*; Supplementary Fig. 9) and woolly mammoth (*Mammuthus primigenius*; Supplementary Fig. 10), detected in four sediment layers sampled (HyL1-3 and EH1), and reindeer (*Rangifer tarandus*; Supplementary Fig. 11) and red deer (*Cervus elaphus*; Supplementary Fig. 12), were restricted to the Hyaena Land samples.

In addition to these, we obtained tentative assignments to other species, including Arctic lemming (*Dicrostonyx torquatus*; Supplementary Figure 13) and otter (*Lutra lutra*; Supplementary Table 7), which had high deamination values (>40%; Supplementary Table 7), but low coverage prevented us from obtaining narrow confidence intervals for deamination. Lower abundance taxa included canids (*Canis lupus/Canis lupus familiaris*) with deamination signatures detected in HyL3, and trout (*Salmo trutta*) in HyL2-3 (Fig. 2A; Supplementary Table 7). In contrast, human (*Homo sapiens*; Supplementary Fig. 14), cattle (*Bos taurus*; Supplementary Fig. 15) and wild boar/pig (*Sus scrofa/Sus scrofa domesticus*; Supplementary Fig. 16) reads consistently showed <20% deamination (Supplementary Table 7), suggesting a more recent origin. Of these species, otter and trout have not been found in Castlepook Cave’s bone assemblages.

We confirmed the species classification results obtained by ancient DNA-aware taxonomic classifiers using blast (Supplementary Fig. 17). While the vast majority of species identified were concordant across methods, these methods also provided erroneous assignments. For example, euka detected Felidae reads which were assigned to *Leptailurus serval* (Supplementary Fig. 18) but these were reassigned to hyena using blast (Supplementary Fig. 19).

Taxa with authentic aDNA patterns were concentrated in Hyaena Land and to some extent in Elephant Hall, with none being detected from the Lower Bone Passage samples. Consistent with these patterns, we observed striking differences in the level of deamination values across the areas, with higher mean 5’-end deamination in Hyaena Land (46.35%) and the Elephant Hall (49.66%) than in the Lower Bone Passage (17.30%) (Figure 2C). In agreement with this, mean fragment length (Figure 2D) was shorter in Hyaena Land (52.51 bp) and Elephant Hall (56.08 bp) than in Lower Bone Passage (66.64 bp). These signals are consistent with the presence of higher amounts of authentic aDNA in Hyaena Land and Elephant Hall than in Lower Bone Passage.

### Phylogenetic analyses

Following species identification in the different areas in Castlepook Cave, we investigated the mitochondrial lineages for which sufficient reads were recovered for informative phylogenetic analyses. Across species, Castlepook Cave sequences fall within previously described contemporary Eurasian mitochondrial diversity.

The cave hyena *(Crocuta crocuta spelaea)* was the most abundant species in our data, providing informative placements for five Castlepook samples, including three from Hyaena Land (HyL1-3) and two from the Elephant Hall (EH1,3). All belonged to mitochondrial haplogroup A1, a lineage distributed in Late Pleistocene European cave hyenas typically older than 37 kya[26], extending the western range of this lineage to Ireland (Figure 3, Supplementary Figure 20). Its sister lineage A2 has been found in pre-LGM Britain[27] and Poland across a wide temporal range (one dating to >44 kya and the other from ∼30 kya)[26], but persists in present-day African spotted hyenas (*Crocuta crocuta*)[28], whereas A1 became extinct in prehistoric times.

**Figure 3.**
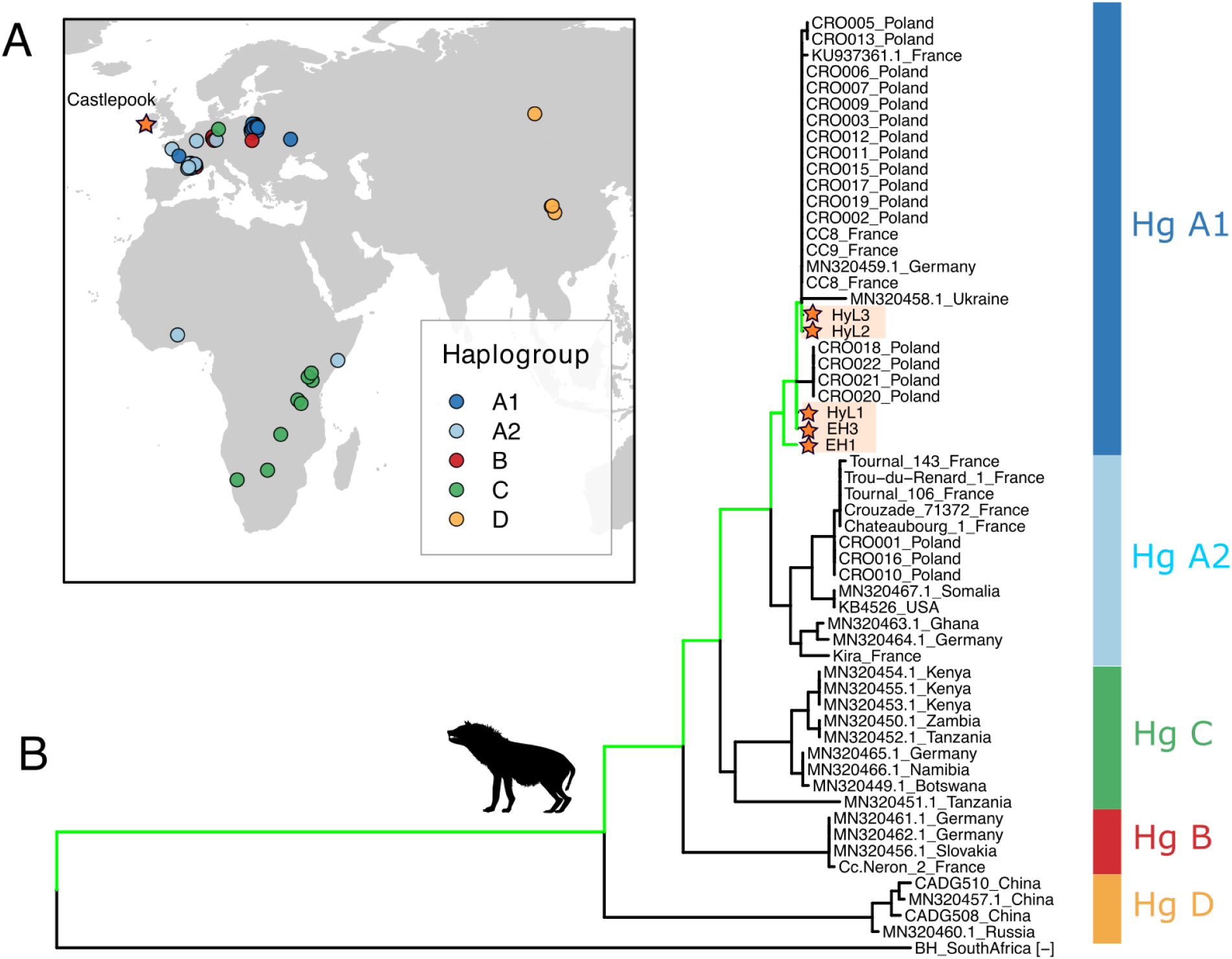
A) Approximate geographical locations of ancient and present-day *Crocuta crocuta and Crocuta c. spelaea* mitochondrial lineages. B) Phylogenetic placement analysis of Castlepook sedimentary aDNA samples into a tree composed of *Crocuta crocuta and Crocuta c. spelaea* mitochondrial lineages and using *Parahyena brunei* as an outgroup.

We detected woolly mammoth DNA (*Mammuthus primigenius*) in four samples, three from Hyaena Land (HyL1-3) and one from the Elephant Hall (EH1), which were placed in the woolly mammoth clade III/B2. This lineage was primarily distributed across western Eurasia prior to ∼30 ka, including in a 48-thousand-year-old Scottish sample[29], and to a lesser extent in Siberia (Figure 4; Supplementary Fig. 21), supporting a pre-LGM age for Castlepook Cave woolly mammoths. Two additional low-coverage samples (EH2-3) were tentatively placed in the branch leading to clade III B1/B2. It is possible that additional data from these two samples would reveal a III/B2 placement given that they carry an ancestral allele at branch III/B1 (Supplementary Fig. 21), but we cannot exclude the possibility that they belong to an unknown clade III lineage. Although these two samples were undetected by the taxonomic classifiers, we confirm that they contain woolly mammoth sequences using blast (Supplementary Fig. 17).

**Figure 4.**
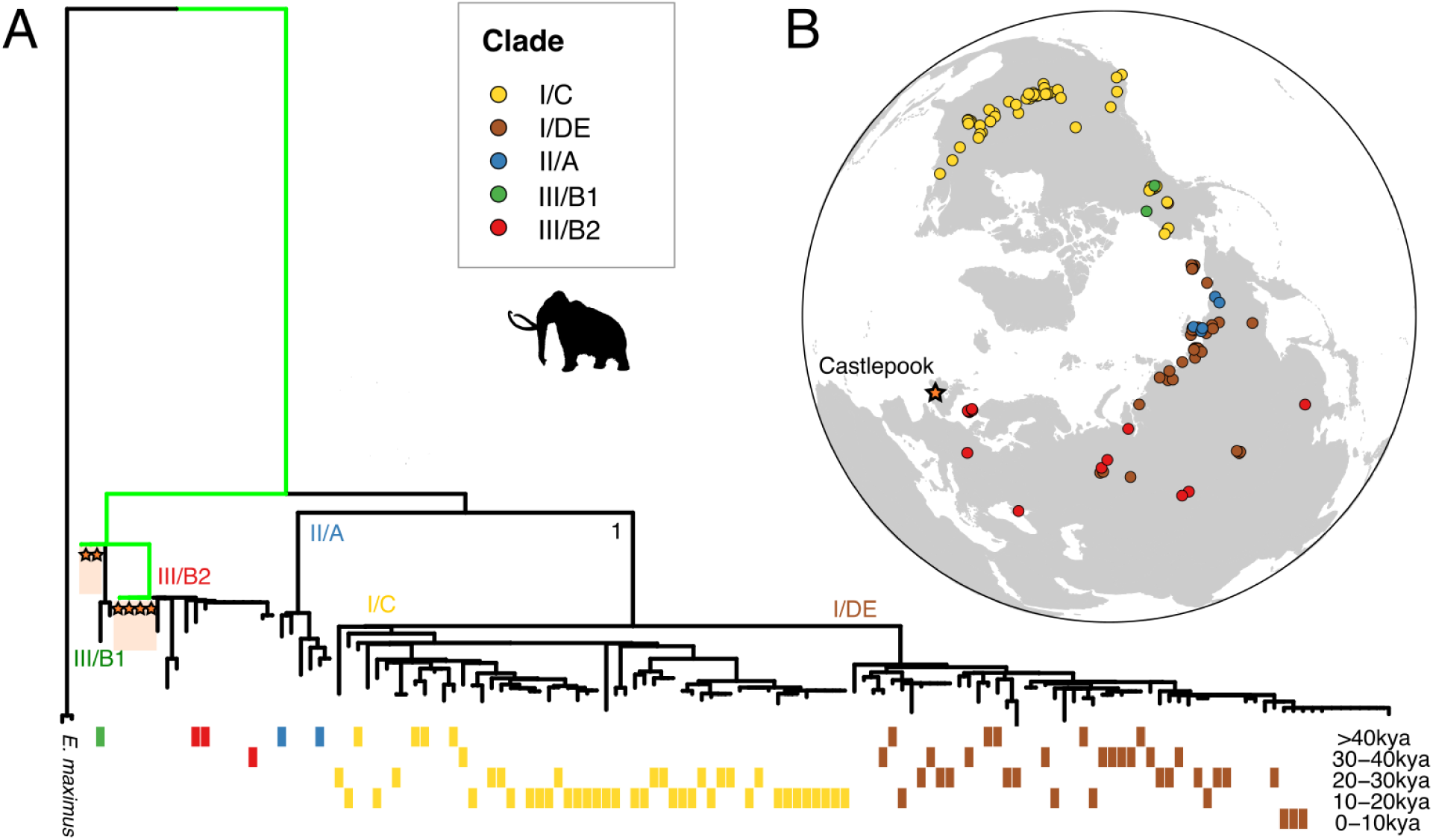
A) Phylogenetic placement analysis of Castlepook Cave sedimentary aDNA samples into a tree composed of mammoth mitochondrial lineages and using elephants as an outgroup. B) Approximate geographical locations of *Mammuthus* mitochondrial lineages.

We detected brown bear (*Ursus arctos*) in all sampled sediments of Hyaena Land and the Elephant Hall (Figure 5), but EH2-3 did not provide sufficient reads for phylogenetic placement. We identified two distinct lineages within four samples (HyL1-3 and EH1): in all four we found clade 1b, which was previously determined to be present in pre-LGM Irish brown bears, and clade 2 in one sample (HyL2; Supplementary Fig. 22), which replaced clade 1b in Ireland at around ∼32 kya and is associated with polar bear introgression [11]. Their co-occurrence in HyL2 suggests either temporal overlap or stratigraphic mixing of sample deposits[30].

**Figure 5.**
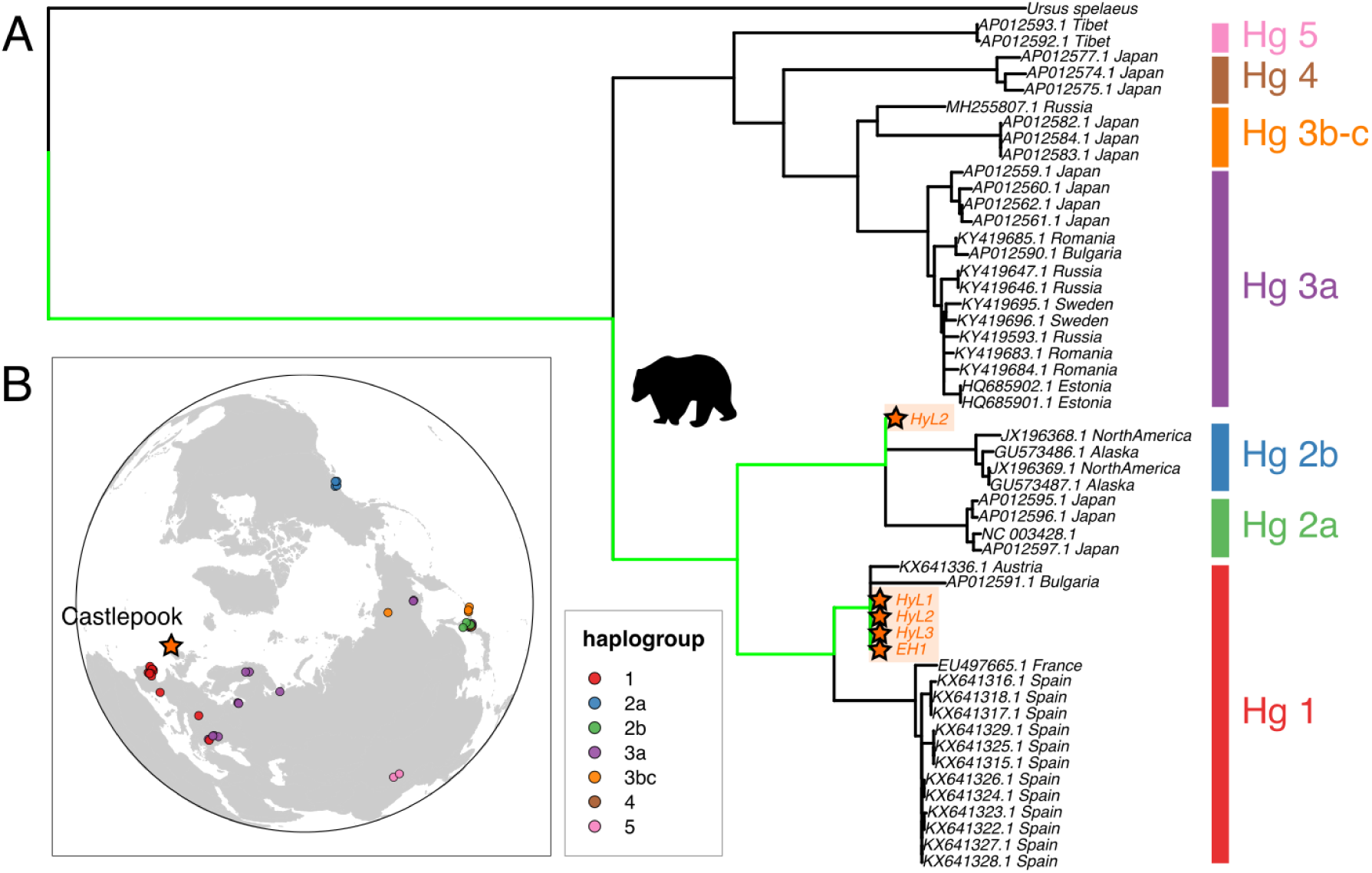
A) Phylogenetic placement analysis of Castlepook sedimentary aDNA samples into a tree composed of bear mitochondrial lineages and using *Ursus spelaeus* as an outgroup. B) Approximate geographical locations of bear mitochondrial lineages.

Ancient genomic data further refined morphological species identifications. Although Arctic fox has previously been reported in Castlepook based on skeletal findings [13], both phylogenetic placement (Supplementary Figure 23) and blast classification (Supplementary Figure 17) of sedaDNA sequences support an assignment to red fox, specifically to the Eurasian clade of mitochondrial diversity (Supplementary Figure 24). Red deer (*Cervus elaphus*) sequences (HyL1-3) belong to the west Eurasian clade A, of western and central European distribution [31] (Supplementary Figure 25).

Hominidae sequences were placed within present-day human mitochondrial diversity (*Homo sapiens,* with one of the samples (HyL1) being tentatively assigned to mtDNA haplogroup V (Supplementary Fig. 26), which occurs at high frequency in present-day Saami [32], whereas most European hunter-gatherers predominantly belonged to haplogroup U [33].

Cattle and wild boar/pig sequences showed limited phylogenetic resolution. The weak signal obtained from cattle reads sequences pointed to *Bos taurus* mtDNA macro-haplogroup T (Supplementary Figure 27). We were unable to distinguish between wild and domesticated mtDNA *Sus scrofa* lineages, with soibean placing the samples at the ancestral branch (Supplementary Figure 16).

In summary, Castlepook Cave faunal sequences consistently fall within West Eurasian diversity, supporting connection to the mainland before the LGM.

## Discussion

Through the analysis of sedimentary ancient DNA from Castlepook cave, we reconstruct Late Pleistocene faunal composition, corroborating and extending the zooarchaeological record, and providing new insights about the population affinities of Irish fauna. Our results are consistent with Ireland forming part of an extended Eurasian population network during the Pleistocene before the LGM, despite Ireland’s peripheral positioning and periods of insularity.

Phylogenetic analyses revealed that mtDNA lineages from Ireland across multiple taxa belong to previously identified haplogroups in continental Europe: cave hyenas belong to haplogroup A1, associated with older Late Pleistocene European cave hyena populations [34], and woolly mammoths belong to clade III/B2, the dominant lineage in Europe and Western Siberia between ∼45.3-31.2 kya, disappearing at around 30 kya. The identification of two distinct brown bear lineages in Castlepook Cave - clade 1b, the predominant pre-LGM lineage in Ireland, and clade 2, which replaced clade 1b in Ireland at around 32 kya - supports a previously-described process of population turnover occurring before the LGM [11].

The pattern of shared mtDNA lineages we recovered is consistent with paleogeographic reconstructions suggesting intermittent connections between Ireland, Britain and continental Europe caused by lower sea levels and glacial expansions during Marine Isotope Stage 3 (55-33 kya)[4]. These land and ice bridges enabled episodic faunal colonisation of Ireland during periods of climatic amelioration, in this way connecting Ireland to a broader European faunal network during the Late Pleistocene.

Our results further constrain the likely timing of these migration processes. The finding of the woolly mammoth lineage III/B2 and the cave hyena haplogroup A1, both associated with pre-LGM European populations, is consistent a pre-LGM source for the corresponding Castlepook Cave faunal assemblages consistent with colonisation during the MIS3. The co-occurence of two distinct bear lineages further suggests that we are capturing a pre-LGM period of population turnover within Ireland, although we cannot exclude processes such as sediment mixing or intrusion occurring between layers[30].

In addition to clarifying population relationships, our results confirm and in some cases improve assessments of faunal composition beyond what can be observed in the bone record. The detection of taxa which are thus far absent from the zooarchaeological record, such as trout and otter, and the genetic identification of red fox rather than Arctic fox as suggested from the animal bones [13], highlights the ability of sedaDNA to recover otherwise unseen species and possibly resolve morphologically ambiguous taxa.

The variation in aDNA preservation across cave contexts - higher in Hyaena Land and the Elephant Hall, and lower in the Lower Bone Passage - highlights the importance of depositional and taphonomic processes in shaping aDNA records [30], and this, in turn, contributes to our understanding of cave formation and sedimentation history, as we can further investigate this preservation difference in the future through other proxies.

Taxa such as human, cattle and wild boar/pig showed low deamination and longer fragment lengths, consistent with more recent origins, rather than deriving from Pleistocene presence within the cave. This observation is consistent with recently obtained radiocarbon dates from charcoal recovered from sediments in the Lower Bone Passage, pointing to a Medieval date, which suggests possible intrusions or deposition into this part of the cave during this period. However, more recent sources for the introduction of contaminant DNA cannot be excluded, including during excavation or exploration of the cave, or during sampling or in the ancient DNA laboratory. Supporting this possibility, our extraction control presented a small proportion of assigned reads to chordates (∼1.5%; Supplementary Table 4), including taxa with deamination (Supplementary Table 7). This signal may reflect minor cross-contamination during lab work, although index hopping during sequencing may also have contributed to this signal. Importantly, the extraction control underwent the same library amplification, target capture and post-capture amplification procedures as the sediment samples, despite contributing only a very small volume to the capture pool, likely leading to an overrepresentation of trace contaminant molecules in the final sequencing data.

Our approach has several limitations. Firstly, we focused on mitochondrial sequences due to their higher abundance in cells and shorter size when compared to the nuclear DNA, making it more cost-effective to target complete mitochondrial sequences from different species. However, due to their nonrecombinant nature and uniparental mode of transmission, mtDNA only provides information about the maternal history of a species. Secondly, to be able to verify ancient DNA authenticity from a small number of unique sequences, we opted against enzymatic treatment of DNA extracts. The post-mortem deamination and short fragment lengths typical of aDNA likely affected taxonomic classification, read alignment and phylogenetic analyses[35]. Despite this, we developed and supported our inferences with multiple methods. Thirdly, the low template nature of sedimentary ancient DNA inevitably leads to extremely high duplication rates (here ∼99%) and consequently to an extremely low number of uniquely mapped reads per species. Fortunately, the mitochondrial DNA is mutation dense [36], and therefore even a small number of reads can substantially improve phylogenetic placement resolution.

In conclusion, our work suggests that Ireland was part of a wider Eurasian faunal network during the Late Pleistocene, consistent with intermittent connectivity driven by sea-level changes and glacial expansions, rather than long term isolation of Irish populations. Our findings also demonstrate the potential of sedaDNA to clarify the genetic affinities of regionally extinct species. Future research, with more extensive sampling of Castlepook Cave and other caves in the region and sequencing nuclear aDNA directly from bones or from sediment using targeted approaches, will enable us to access more finely resolved information about the ancestry, timings of colonization and admixture events, and demographic responses of ancient megafauna to Pleistocene environmental change.

## Methods

### Sampling of sediment samples from Castlepook Cave

We collected sediment samples from three Castlepook Cave areas (Hyaena Land, the Elephant Hall and the Lower Bone Passage) in June 2022 and September 2024. We sampled an area within each context by collecting sediments using a sterile scalpel and storing these in 50 mL falcon tubes. Where possible, sediments were collected ∼10-20 cm into the excavation trench wall, away from the exposed face of the sediment profile, to prevent surface contamination[37].

### Ancient DNA extraction

All ancient DNA work was conducted at LJMU’s Ancient DNA Lab. We extracted approximately 5 g from a total of eight samples from sediment layers in three different areas in Castlepook Cave (Supplementary Table 1) using a modified protocol[38] to account for the larger amount of sediment extracted in the present work in comparison to the original protocol[39] (50 mg). We weighed 5 g of each sample and placed it into sterile 50 ml tubes, including an extra empty tube as a full process control.

We added 5 ml of an upscaled lysis buffer[39] to each of the nine tubes, including 4.5 ml of 0.5M EDTA (pH 8.0), 2.5 µl of Tween 20, 125µl of 10 mg/ml proteinase K and 372 µl of nuclease-free water, mixing the samples by vortexing for ten seconds and by gently flicking and inverting to prevent the formation of a sediment pellet. We irradiated the lysis buffer for 30 minutes under UV light before adding the proteinase K. We incubated the samples at 37°C for 24 hours, rotating at 18 rpm to ensure thorough mixing. Following incubation, we concentrated the samples through sequential centrifugation and decantation, resulting in a progressively smaller volume, until 500 µl of clear lysate remained, as previously described[38].

We purified the lysates by adding 5.2 ml of extraction buffer D (5M guanidine hydrochloride, 40% 2-propanol, 0.12M sodium acetate and 0.05% Tween 20) to 500 µl of lysate in preassembled silica spin columns and collection tubes (High Pure Viral Nucleic Acid Large Volume Kit) and mixed by pipetting and inverting[39]. The remaining extraction purification steps were completed following the original protocol[39].

### Library preparation and PCR amplification

We prepared DNA libraries from 21.25 ul of DNA extracts using modified[40] double-stranded DNA library preparation protocol[41]. We amplified the resulting libraries using AmpliTaq Gold because of its uracyl proofreading ability, using a double indexing strategy with a unique index combination (NEBNext® Multiplex Oligos for Illumina) for each sample and including a PCR control. In order to increase the library diversity, we carried out three PCR reactions with a different index combination per sample using AmpliTaq Gold DNA polymerase for 12 cycles as previously described[42].

We purified the PCR products using the Minelute PCR Purification kit following the manufacturer’s protocol with the exception of eluting in 12 µl of EBT which was pre-heated to 65°C and transferring to new PCR tubes. We quantified the amplified libraries with Qubit (High Sensitivity DNA kit) and visualised fragment size distribution using Tapestation (High Sensitivity DNA kit).

### Mitochondrial DNA enrichment and sequencing

We designed a custom target capture array with Twist Bioscience including mitochondrial sequences belonging to 30 species known or hypothesised to have inhabited Ireland in prehistoric times (Supplementary Table 2).

We pooled amplified libraries equimolarly, including three PCR reactions per sample and two samples per capture reaction, making a total of 6 amplified libraries per capture reaction, and four capture reactions in total. We spiked in 0.1 ul of the PCR and extraction controls into the fourth reaction. We enriched our amplified libraries by following the manufacturer’s protocol. We amplified post-capture reactions using 2.5 µl of primer mix, 25 µl Equinox Library Amplification Mix and 22.5 µl of each reaction for 23 cycles. We purified post capture PCR reactions as recommended in the protocol and quantified the results using Qubit and Tapestation as described above. We pooled PCR reactions equimolarly and sequenced them in one 10B lane of Illumina NovaSeq X at Macrogen Europe, resulting in approximately 1.18 billion paired-end reads.

### Initial data processing and filtering

We trimmed Illumina sequencing reads using AdapterRemoval v. 2.3.2[43] using parameters -minlength 30 --trimns --trimqualities --minquality 2 --collapse. We removed reads with base quality lower than 30 in at least 25% of each read using FASTX-Toolkit v. 0.0.14 and read duplicates with SGA v. 0.10.15 preprocess using a dust threshold of 1, followed by SGA index -a ropebwt and SGA filter with QualTrim 15, requiring a minimum length of 30 bases (https://github.com/jts/sga).

### Taxonomic classification and evaluation of authenticity

We used several approaches for taxonomic classification. First, we used Taxonomy classification of reads using Kraken2[19] and the full nt database, downloaded from https://genome-idx.s3.amazonaws.com/kraken/k2_nt_20240530.tar.gz, and visualised the results using pavian v. 1.0[44]. Kraken2 is not aDNA-aware and produced a few spurious matches. To improve this, we used two aDNA-aware taxonomic classification approaches: the vegan euka/soibean workflow[23,24] and quicksand[22]. Both methods were designed to handle aDNA sequences and prevent spurious assignments by imposing various criteria to establish authenticity, including the evenness or breadth of mitochondrial DNA sequence coverage and deamination, and providing other authenticity metrics such as fragment length.

For classification with vegan v. 3.0.0 euka[23], we used default parameters (minimum of 100 reads assigned to a given taxon, and minimum bin size 6 to ensure coverage evenness) and the provided database that includes 335 arthropodic and tetrapodic taxa. To obtain species-level information from our samples, we used soibean[24] with default parameters to conduct a variant graph approach[45] to species identification directly from the reads assigned by euka to each family. For classification with quicksand v. 2.5, we used default parameters on the raw trimmed reads (because quicksand uses KrakenUnique and it also removes duplicates, whereas euka/soibean require deduplication before analysis) and the RefSeq release v. 221 database. Quicksand’s ancientness score ++ indicates the presence of deamination in >9.5% of the reads in both the 5’- and 3’-ends, whereas with + this signal is only present in one of the ends.

To confirm taxonomic classifications, we used blastn v. 2.16.0+[46] using the nt database on the set of filtered reads. We conducted an LCA analysis of the blast output using MEGAN v. 6.25.10[47], requiring a minimum percent identity of 90 and a minimum of 100 reads assigned to a given taxon, as previously described[48].

### Phylogenetic placement

We used pathPhynder v. 1.2.3[35] for phylogenetic placement of sedimentary ancient DNA samples, which requires a VCF file and a phylogenetic tree for each separate analysis. To prepare a VCF, we compiled a list of accessions of mitochondrial fasta files from previously published samples belonging to species of interest for building a phylogeny, including hyena[26,28,49–53], woolly mammoth, bear[54–63], hominid[64] and deer[31,65].

When necessary, fasta files were concatenated into a multifasta file and aligned using muscle v.5.3[66] using default parameters. We called SNPs from multifasta files using snp-sites[67] and used bcftools v.1.19[68] view with parameters ‘view -m2 -M2 -c1’, to select biallelic variants observed at least in one individual. The proportion of the mitochondrial DNA sequenced in each accession is variable, and in cases where many missing sites are present this may reduce the number of informative variants retrieved from the multifasta file. On the other hand, removing accessions from the multifasta file will result in the loss of informative variants for phylogenetic analysis. Therefore, we aimed to remove low coverage fasta sequences while maximizing the number of informative sites for each species through an iterative procedure. First, we impose a threshold of the number of missing sites (Ns) to filter individual sequences on the multifasta file (threshold range from 2000 to 500 missing sites in intervals of 50) and use snp-sites to call variants. We then selected the missing sites threshold which recovered the greatest number of variants for phylogenetic analysis

If a phylogenetic tree was not available, we built one using the newly generated vcf file using plink[69] to estimate genetic distance between the samples (--distance square 1-ibs) and the ‘nj’ function of the R[70] package phytools v.1.2-0[71], and rerooted the phylogeny at the midpoint of the longest branch. We then ran phynder to place SNPs on the branches of the phylogeny and pathPhynder ‘prepare’ to prepare tree files for placement.

In order to place the samples into the phylogeny the ancient reads must be mapped onto the same coordinates as the multifasta alignment. To do this, we prepared a consensus sequence of the multifasta file using EMBOSS cons tool v.6.6.0.0[72] with datafile EDNAFULL. We then aligned collapsed fastq files to this consensus reference sequence using bwa aln[73] v.0.7.17-r1198 using recommended parameters that balance sensitivity and accuracy in ancient DNA alignment[45], filtered reads using samtools v. 1.13[74] for mapping quality ≥30 and removed duplicates using sambamba v. 0.6.6[75] markdup.

We placed the ancient DNA samples using pathPhynder’s best path approach with default parameters. Sedimentary aDNA samples may contain mixtures of different species and lineages, leading to the presence of both support and conflict alleles on the same branches and preventing an informative placement. To enable a more downstream placement in cases of high conflict SNPs, we set the -t parameter to a large value (e.g. 100), in this way increasing the number of conflict SNPs tolerated prior to stopping traversing a given path.

In order to be able to disentangle the two distinct bear lineages, namely the brown bear-related clade 1b and the polar bear-related clade 2, we followed a strategy of competitive mapping previously used for black and brown bears[76]. Briefly, we aligned sequencing reads to a multifasta composed of all RefSeq mtDNA sequences with bwa and filtered them as described above. The sequences aligned to the brown bear mtDNA sequence were used to identify clade 1b, whereas the reads mapping to polar bear were used to determine clade 2.

All plots were generated in R[77]. Silhouettes for the different species were obtained from PhyloPic (https://www.phylopic.org/).

## Authors’ contributions

RM: methodology, investigation, formal analysis, data curation, visualization, writing-original draft, writing-review and editing;

STW: investigation, formal analysis, visualization, writing-original draft, writing-review and editing;

TM: investigation;

PK: visualization, resources, writing-review and editing;

RPJ: conceptualization, visualization, resources, supervision, project administration, funding acquisition, writing-review and editing;

RFC: conceptualization, supervision, writing-original draft, writing-review and editing;

ADM: conceptualization, supervision, writing-original draft, writing-review and editing, funding acquisition;

HL: writing-original draft, writing-review and editing, funding acquisition.

All authors gave final approval for publication and agreed to be held accountable for the work performed therein.

## Acknowledgments

We thank Andrew Foote and Jan Laine for advice related to sedimentary ancient DNA extraction, Maciej Krajcarz for assistance with cave hyena metadata. The authors would also like to acknowledge LJMU’s Prospero High Performance Computing team for the provision of computational resources. Samples were taken by members of the Castlepook Cave Project and Irish Caves Bones Project. We thank members of the Cork Speleological Group and the Irish Cave Rescue Organisation for safety risk assessments and excavation support. We thank Mrs Ellen Howell, Mr Sylvester Boland and Mr John-Paul O’Brien for permission for cave access/use of land. We thank the Doneraile community for their ongoing local support, in particular Mr Pat McInearney and Mr Donal Broderick. Thanks to National Monuments Service and the National Museum of Ireland for relevant licences.

## Data availability

Raw sequencing reads were uploaded to the European Nucleotide Archive, accession number PRJEB106894, and will be made available upon publication.

