## Supplementary Information for "Sedimentary ancient DNA reveals Late Pleistocene faunal connectivity between Ireland and Eurasia"

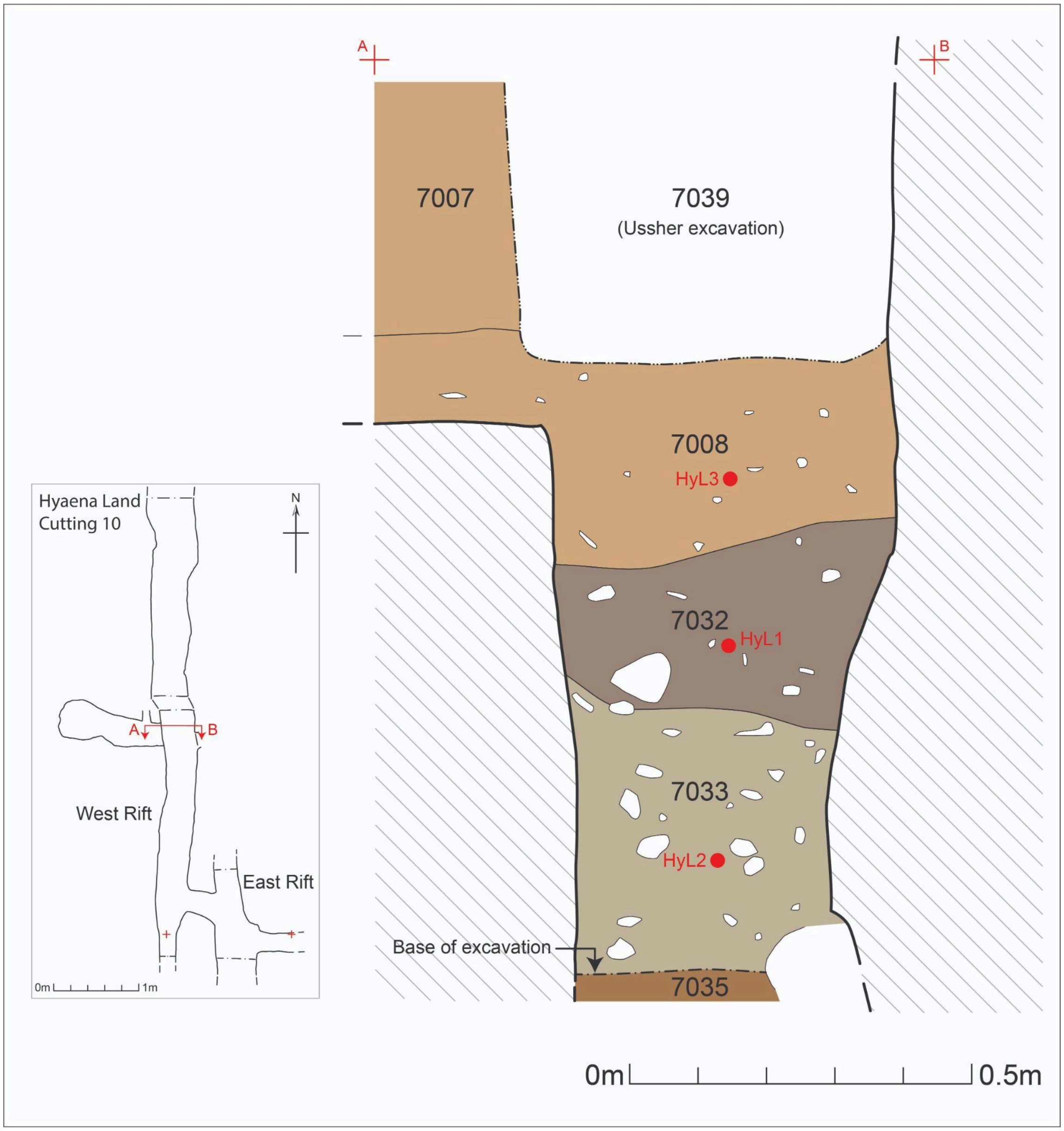


**Supplementary Figure 1** - Section drawing of Castlepook Cave's Hyaena Land context and sediment sample location.

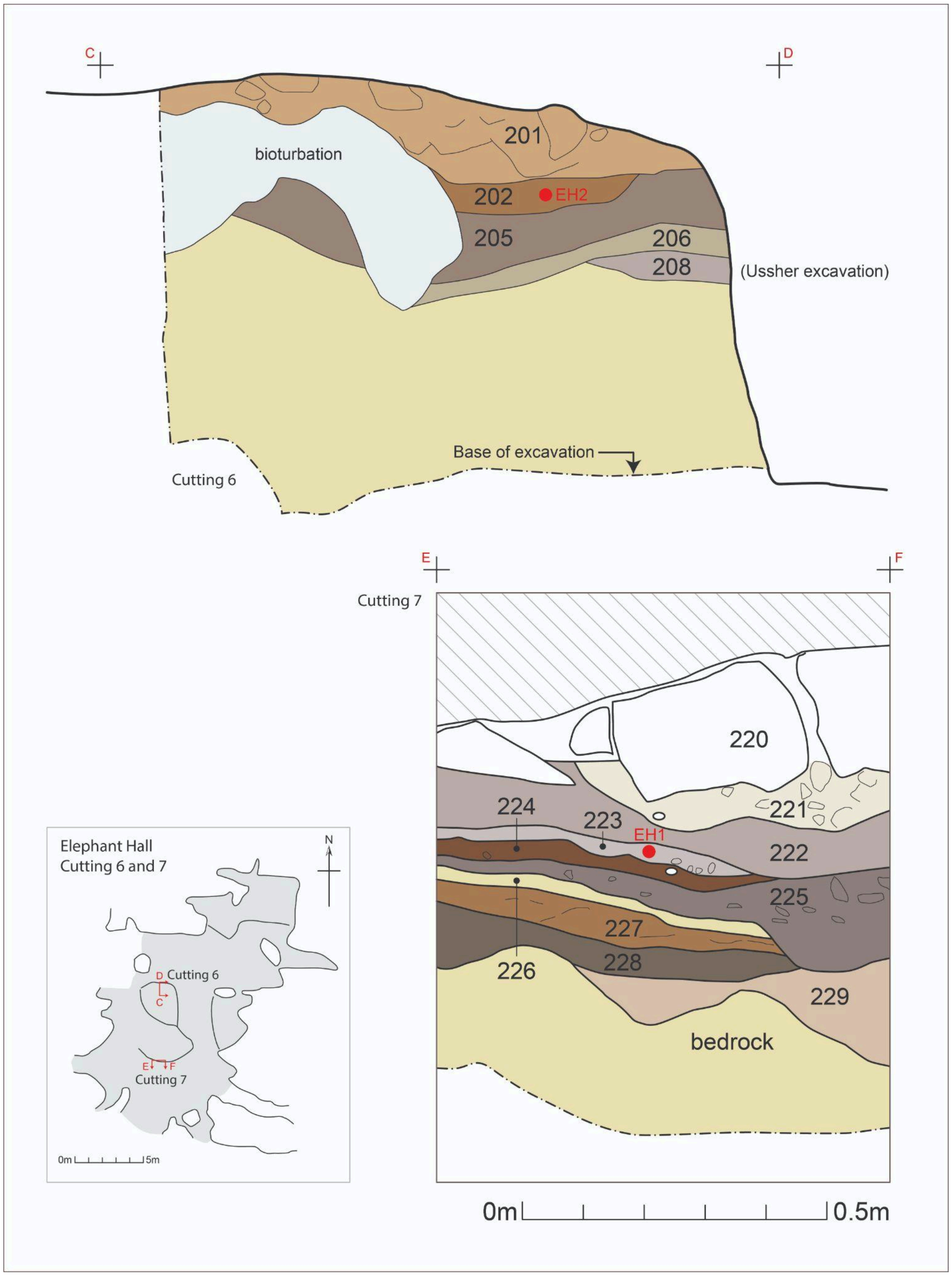


**Supplementary Figure 2 -** Section drawing of Castlepook Cave's Elephant Hall context and sediment sample location.

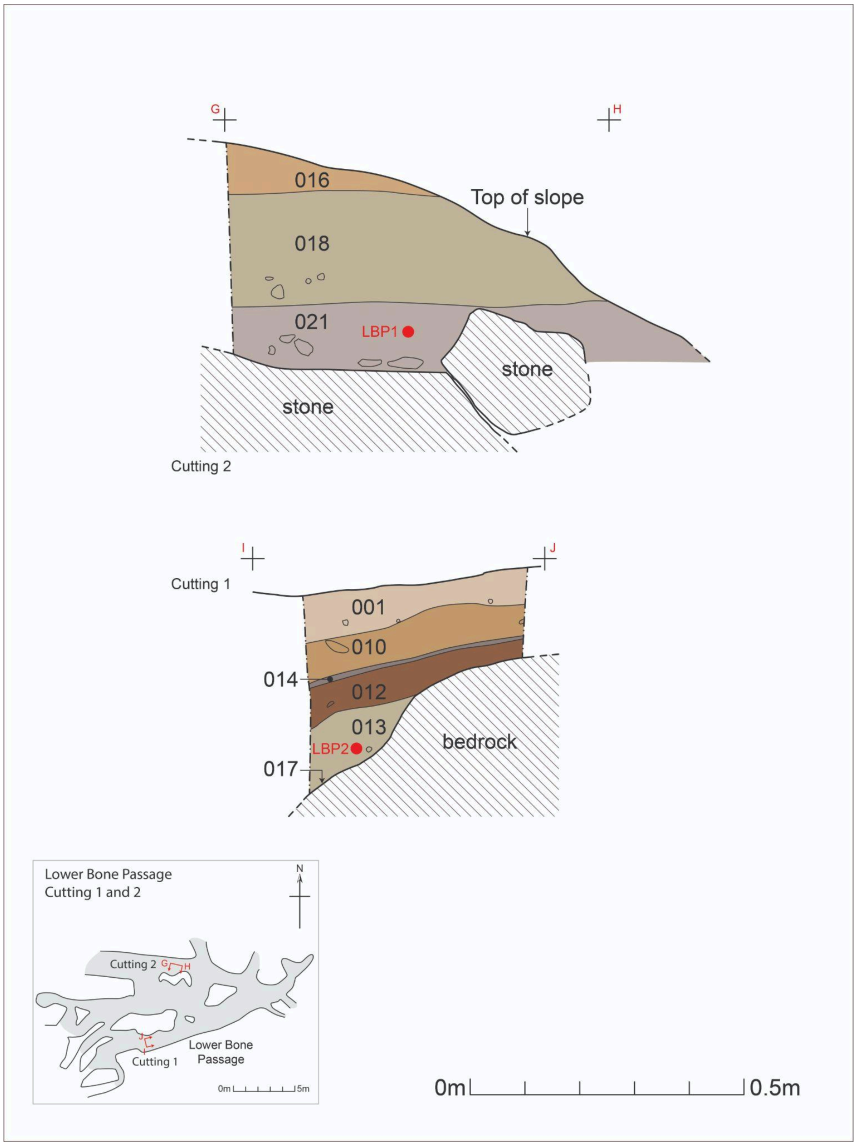


**Supplementary Figure 3** - Section drawing of Castlepook Cave's Lower Bone Passage context and sediment sample locations.

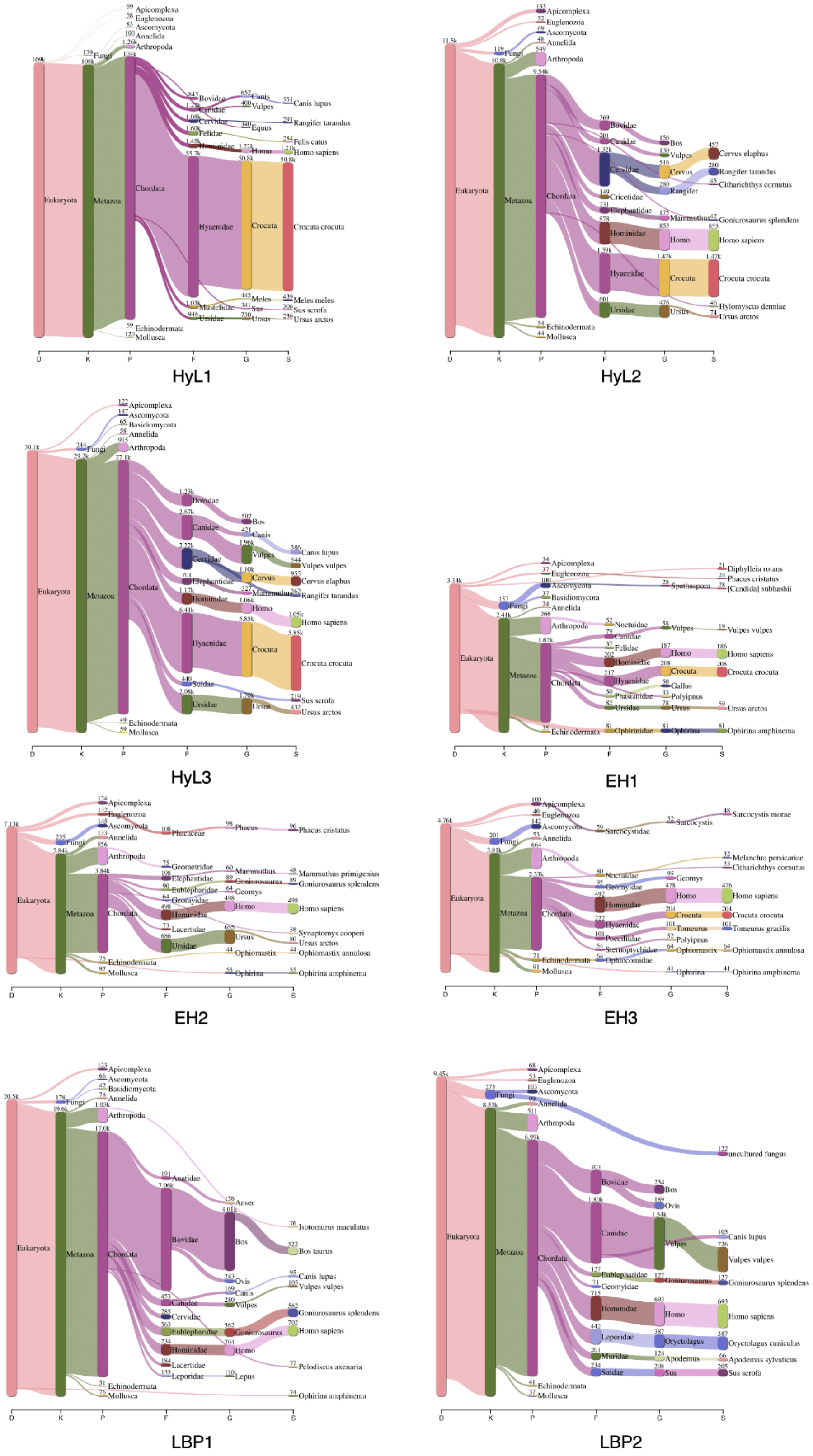


**Supplementary Figure 4** - Pavian visualisation of taxonomic classification results of Castlepook Cave sedimentary ancient DNA sequences.

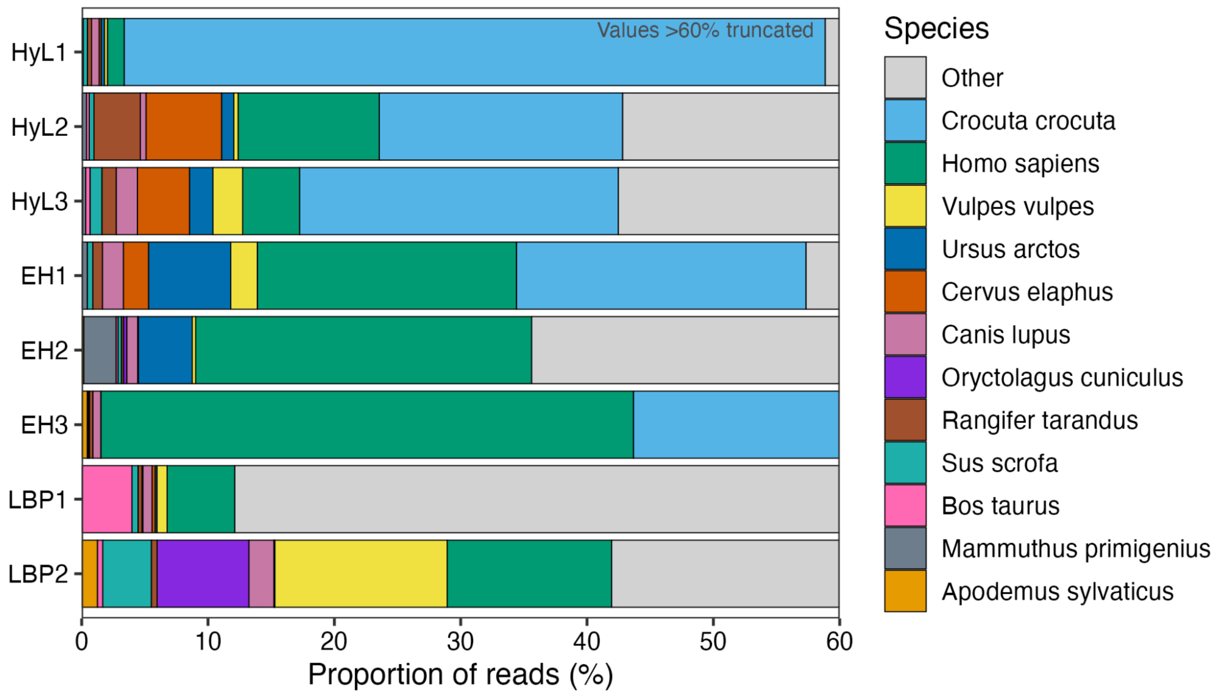


**Supplementary Figure 5** - Kraken classification of mammalian reads.

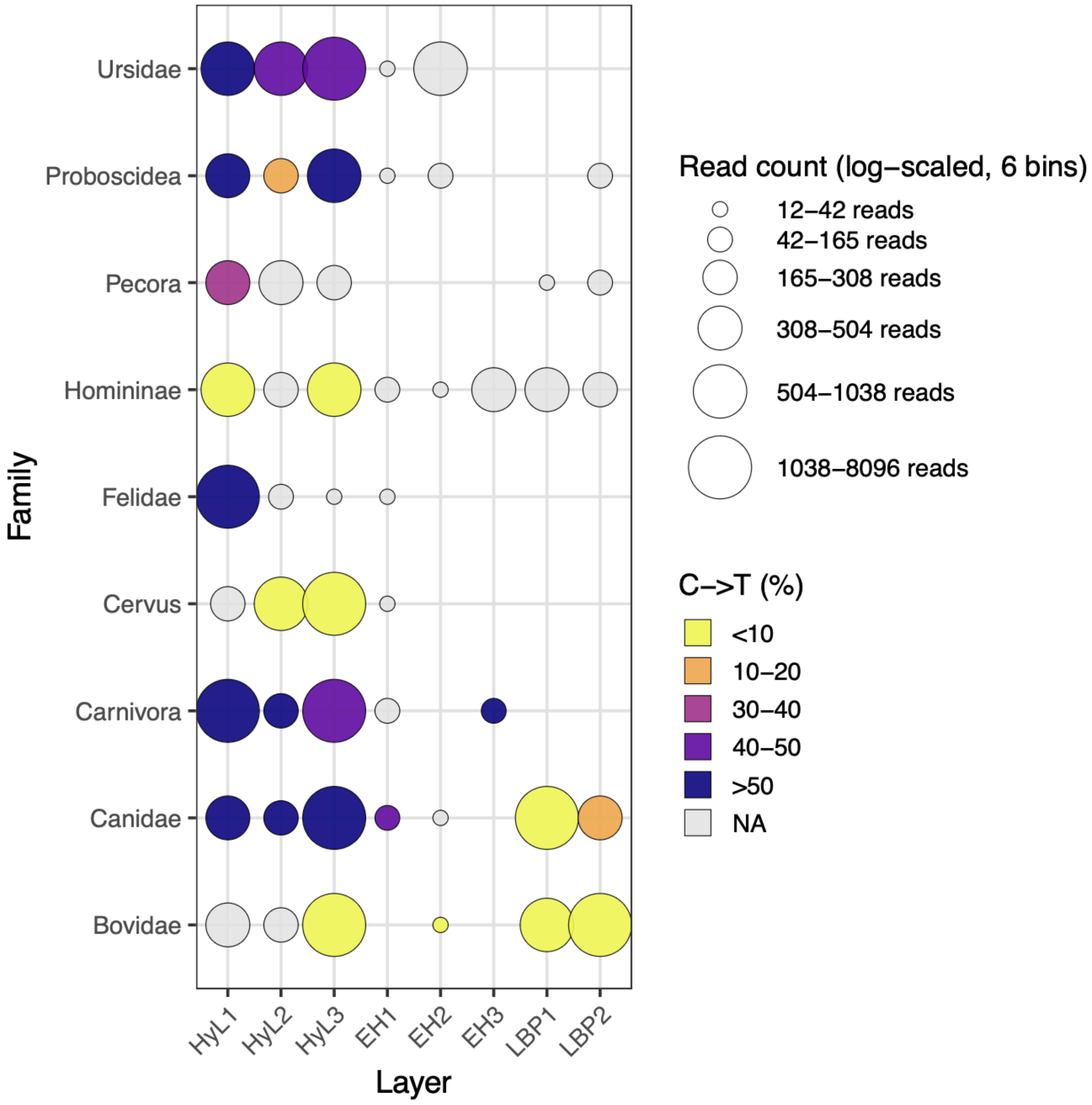


**Supplementary Figure 6** - Taxa abundance in the eight Castlepook Cave layers estimated with euka. Circle size indicates the number of reads assigned to each family (log-scaled) and the colours indicate the frequency of C to T changes (deamination) at the first base for each taxa.

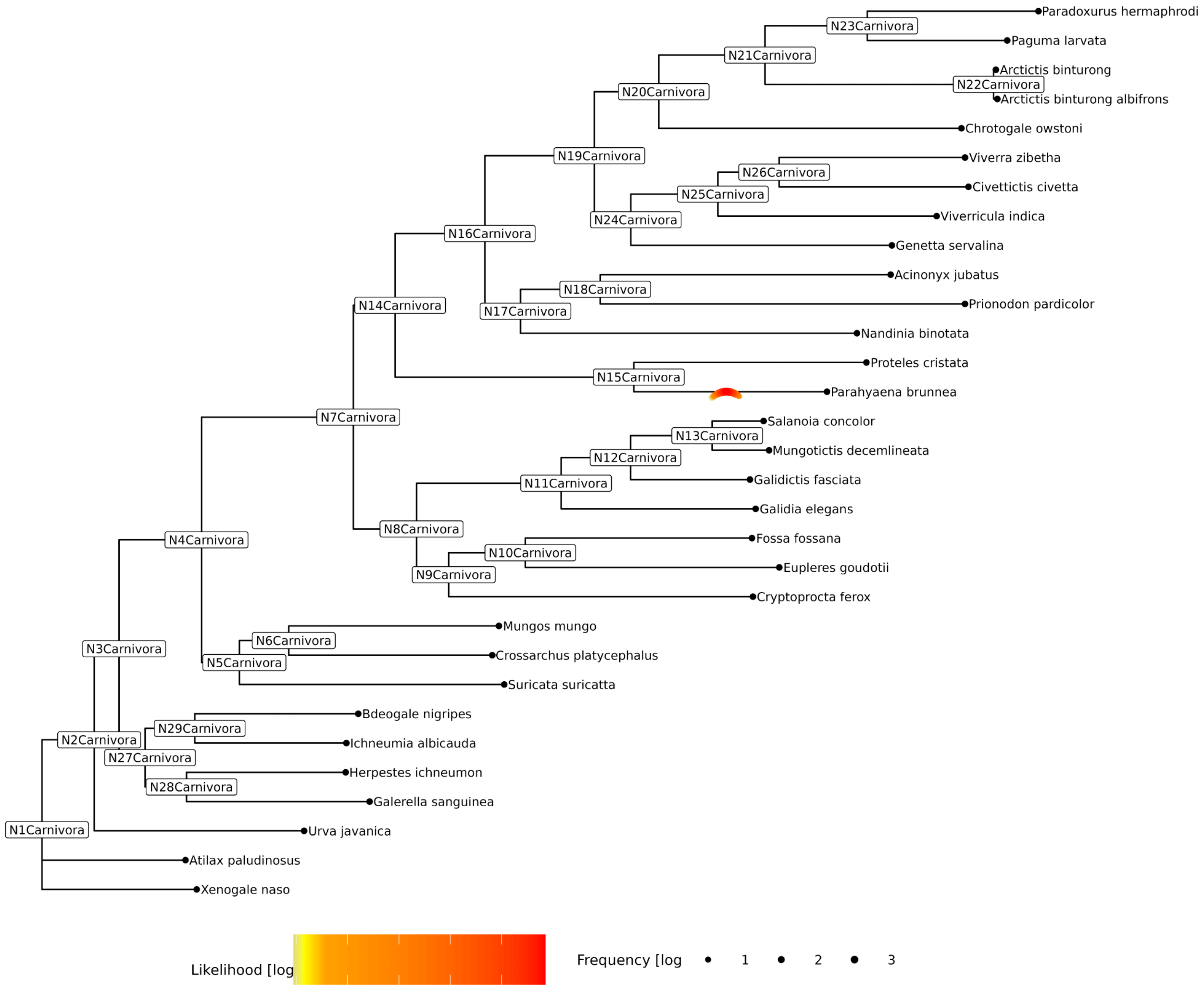


**Supplementary Figure 7 -** Soibean assignment of sequences to *Parahyaena brunnea* (brown hyena) within the Carnivora mtDNA phylogenetic tree. Note that neither Crocuta crocuta or Crocuta crocuta spelaea are present in this tree and therefore Parahyaena brunnea represents the closest representative.

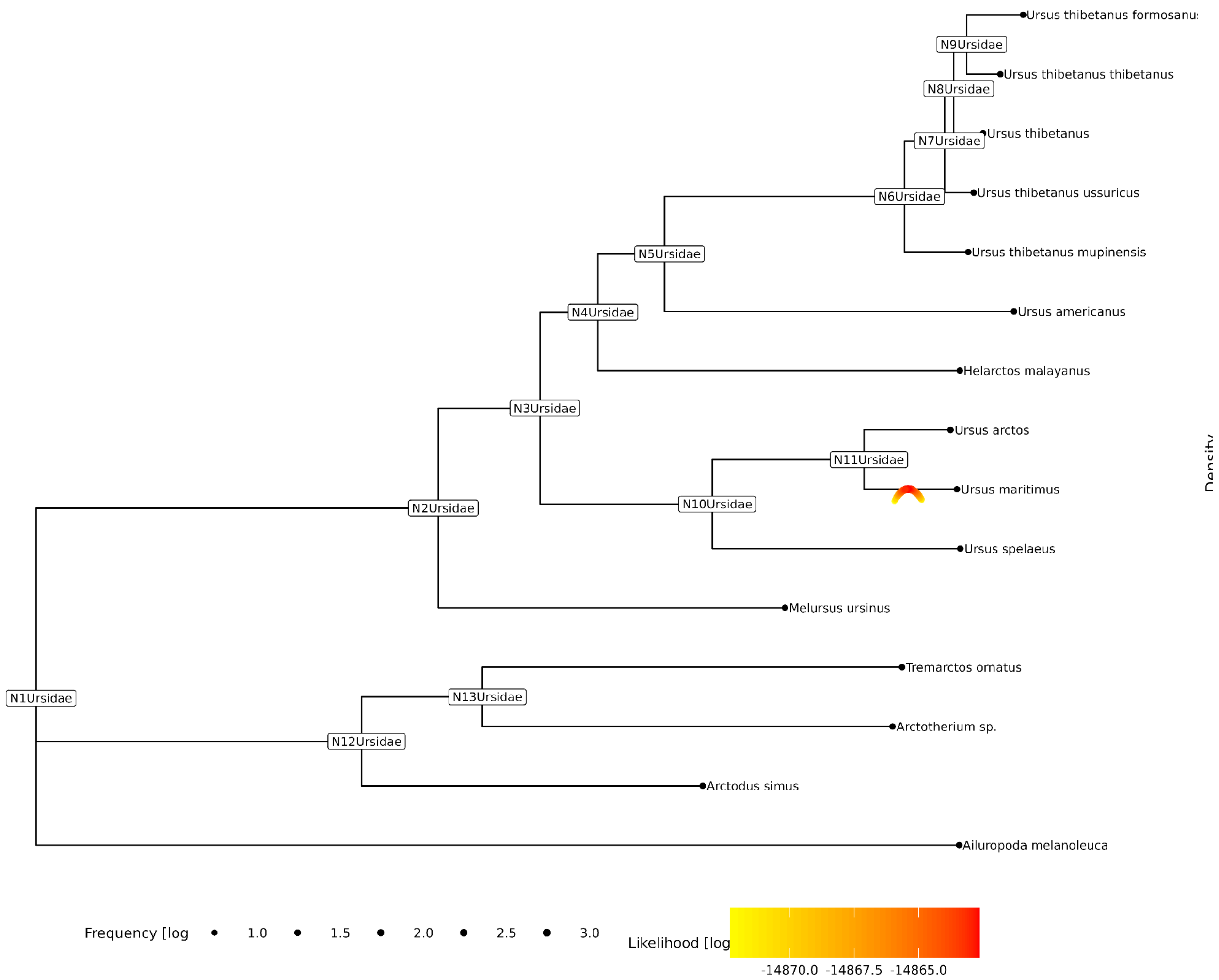


**Supplementary Figure 8 -** Soibean assignment of sequences to *Ursus maritimus* (polar bear) within the Ursidae mtDNA phylogenetic tree. Note that there was an influx of polar bear-derived mitochondrial lineages into Irish brown bears.

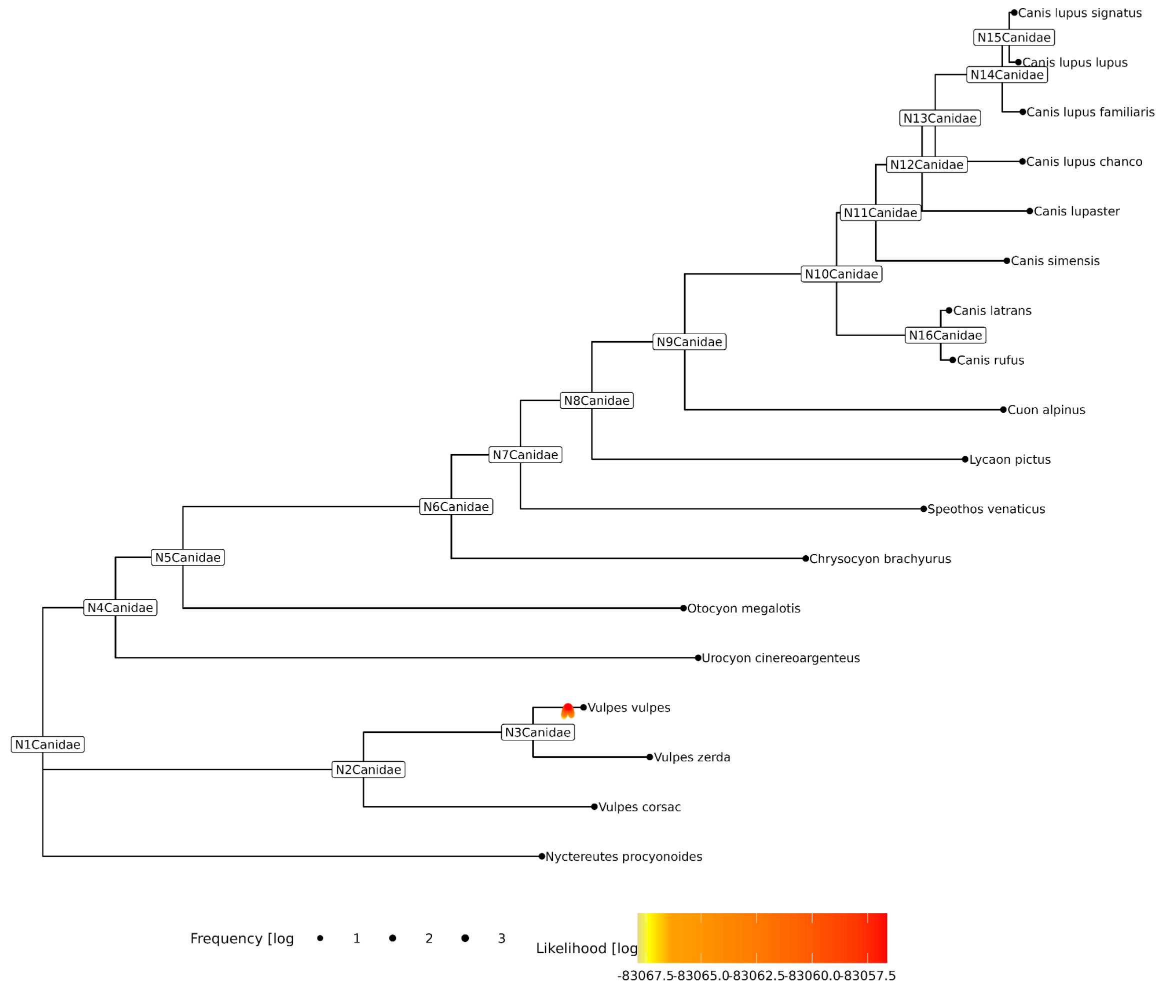


**Supplementary Figure 9 -** Soibean assignment of sequences to *Vulpes vulpes* (red fox) within the Canidae mtDNA phylogenetic tree.

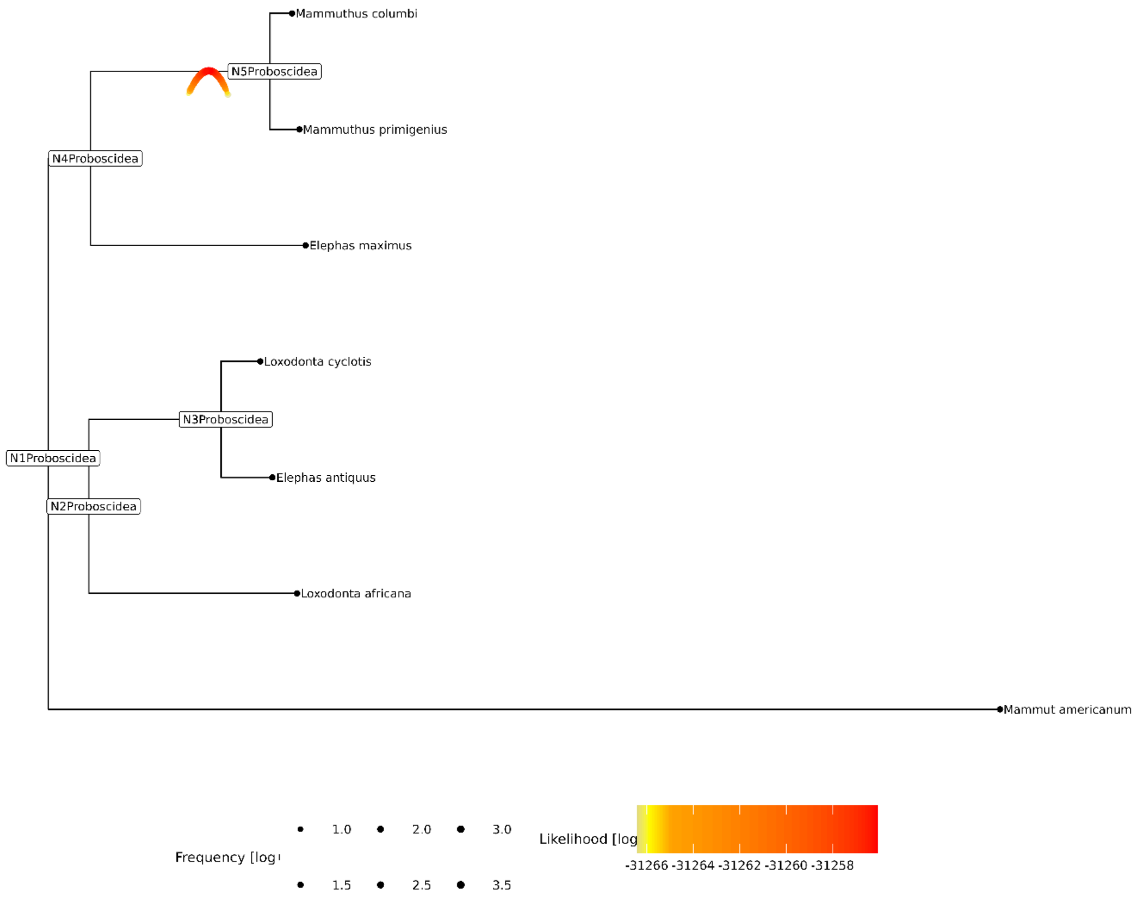


**Supplementary Figure 10** - Soibean assignment of sequences to *Mammuthus* (mammoth) within the Proboscidea mtDNA phylogenetic tree.

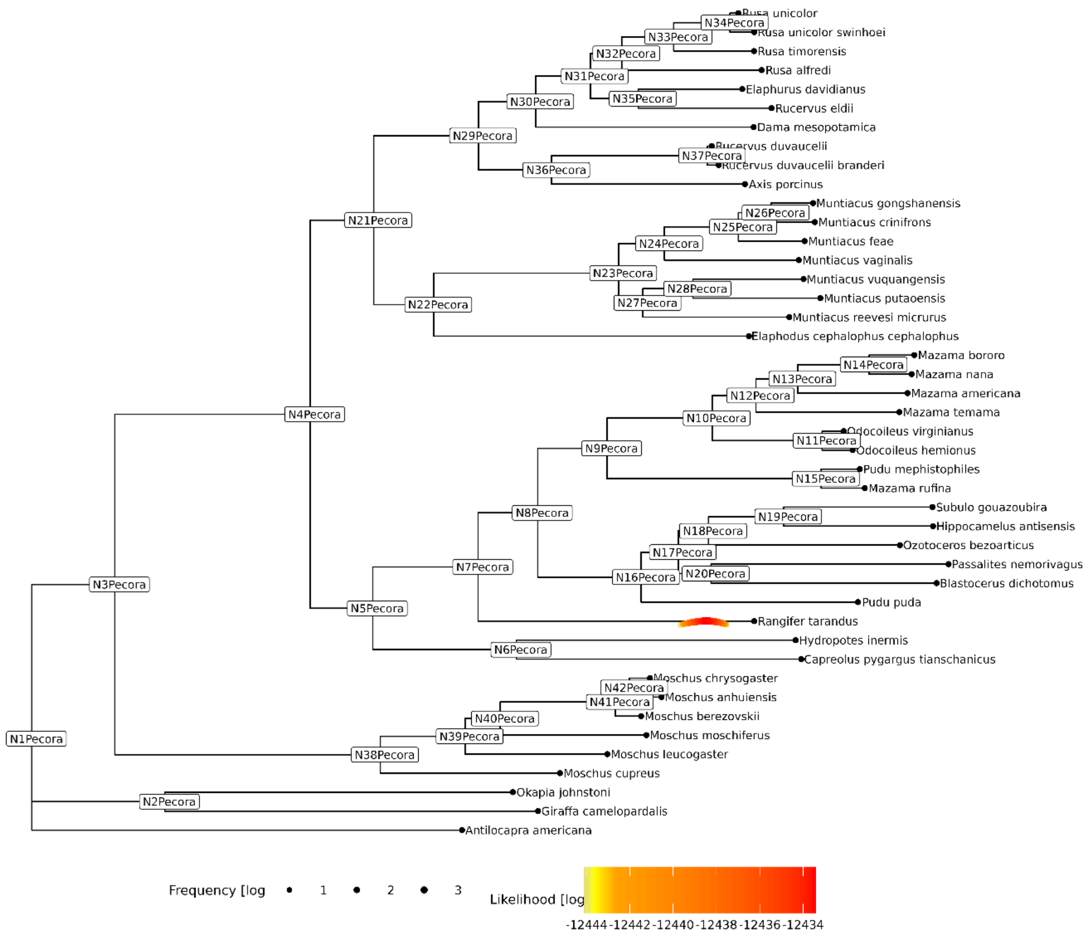


**Supplementary Figure 11** - Soibean assignment of sequences to *Rangifer tarandus* (reindeer) within the Pecora mtDNA phylogenetic tree.

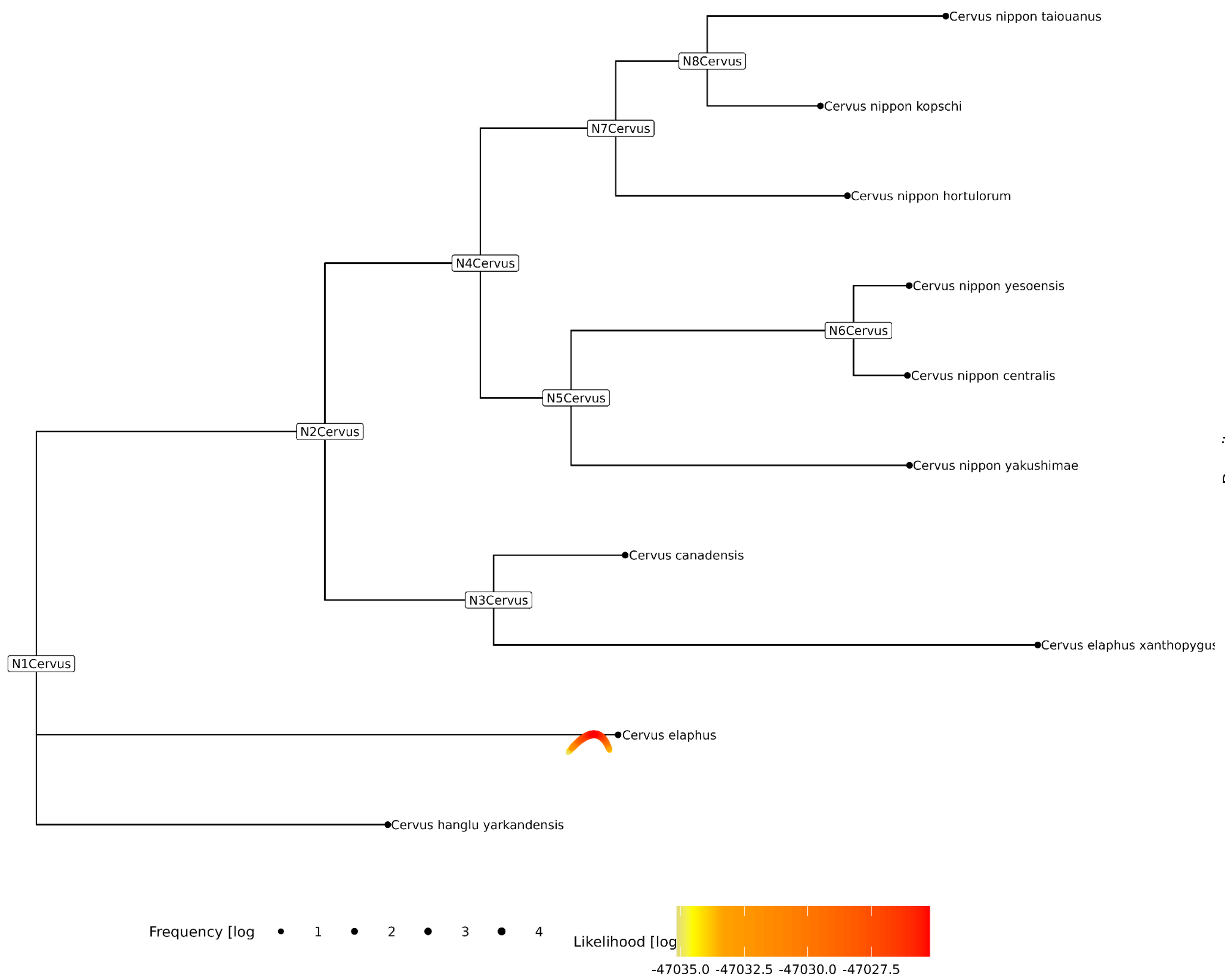


**Supplementary Figure 12** - Soibean assignment of sequences to *Cervus elaphus* (red deer) within the Cervus mtDNA phylogenetic tree.

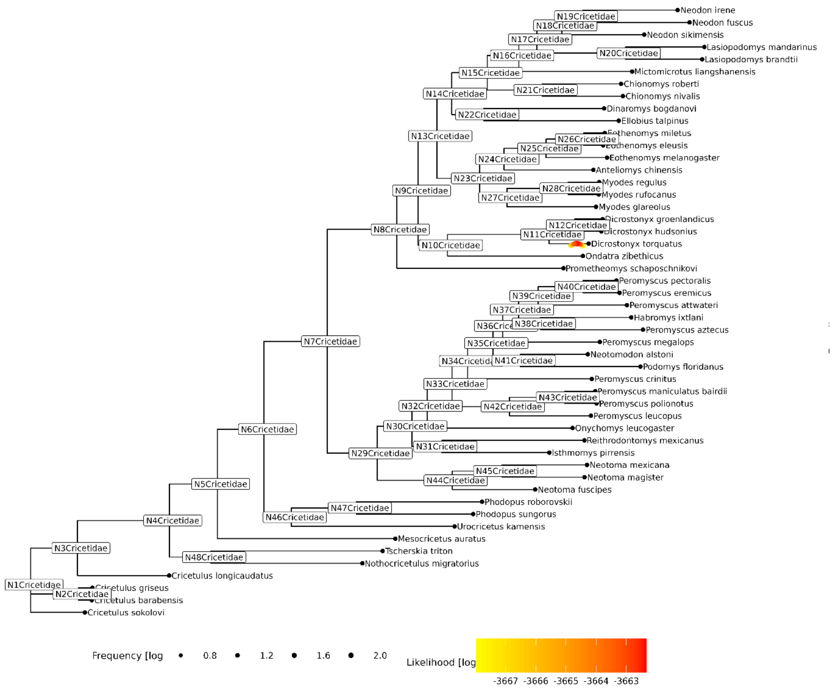


Supplementary Figure 13 - Soibean assignment of sequences to *Dicrotonyx torquatus* (arctic lemming). We confirmed the assignment of these reads using blast.

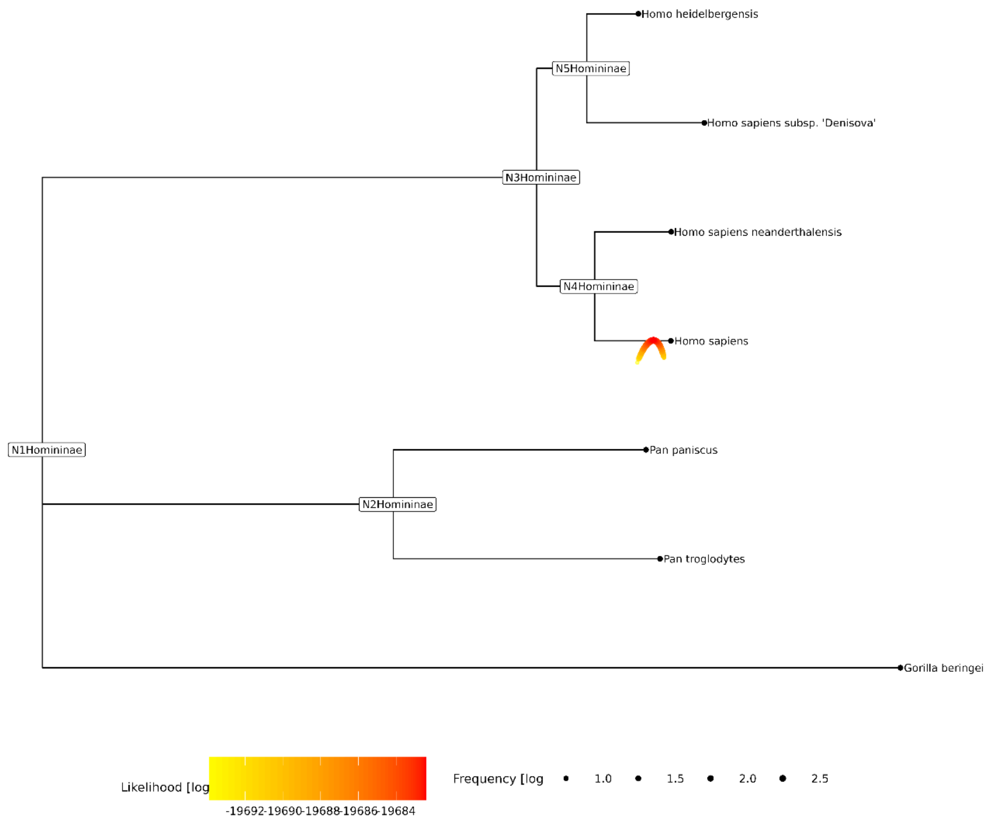


**Supplementary Figure 14** - Soibean assignment of sequences to *Homo sapiens* (human) within the Hominidae mtDNA phylogenetic tree.

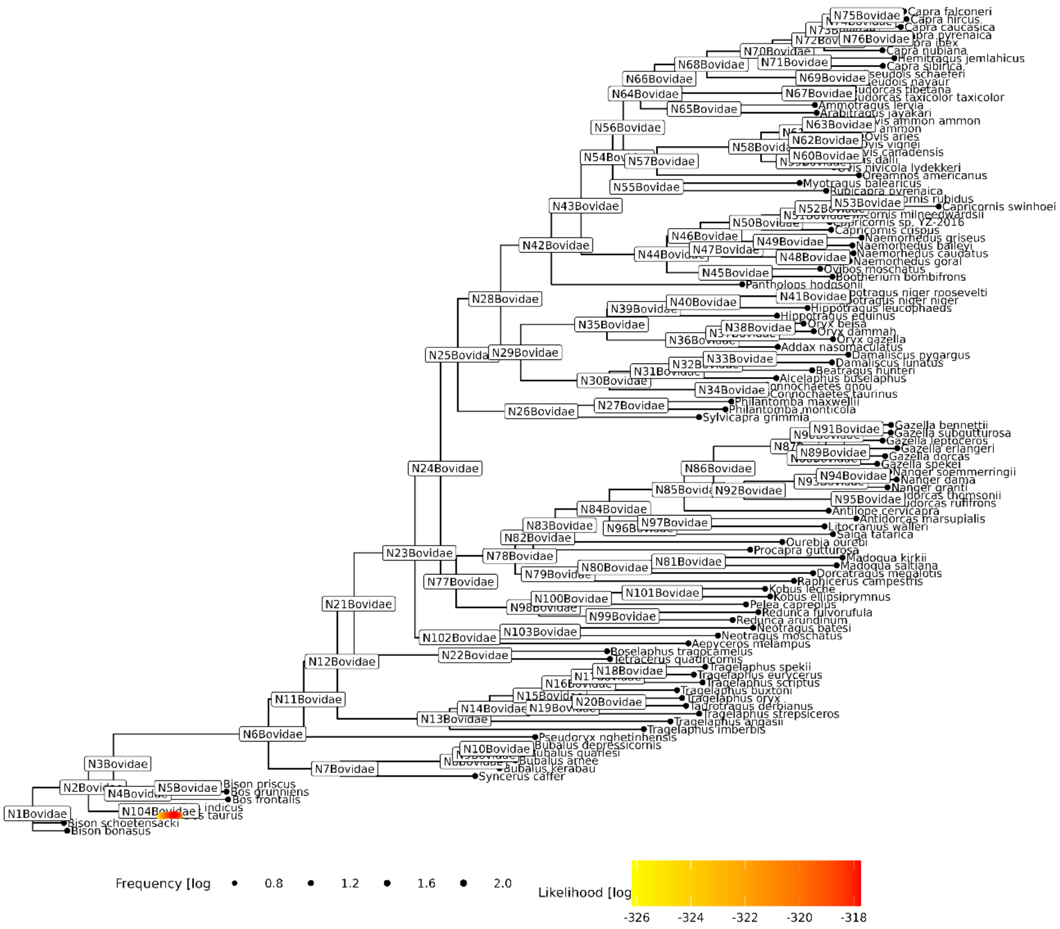


**Supplementary Figure 15 -** Soibean assignment of sequences to *Bos taurus* (cattle) within the Bovidae mtDNA phylogenetic tree.

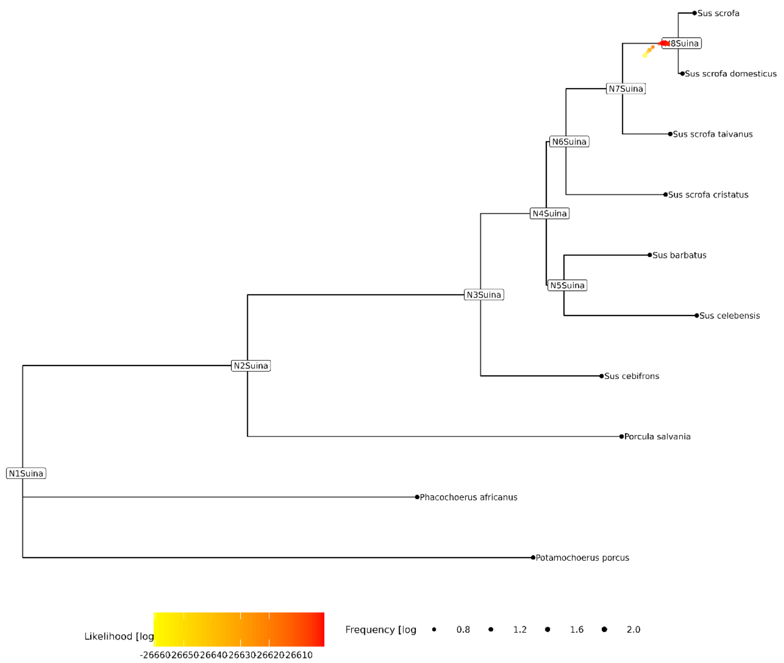


**Supplementary Figure 16 -** Soibean assignment of sequences to *Sus scrofa/Sus scrofa domesticus* (wild boar/pig) within the Suina mtDNA phylogenetic tree. We confirmed the assignment of these reads using blast.

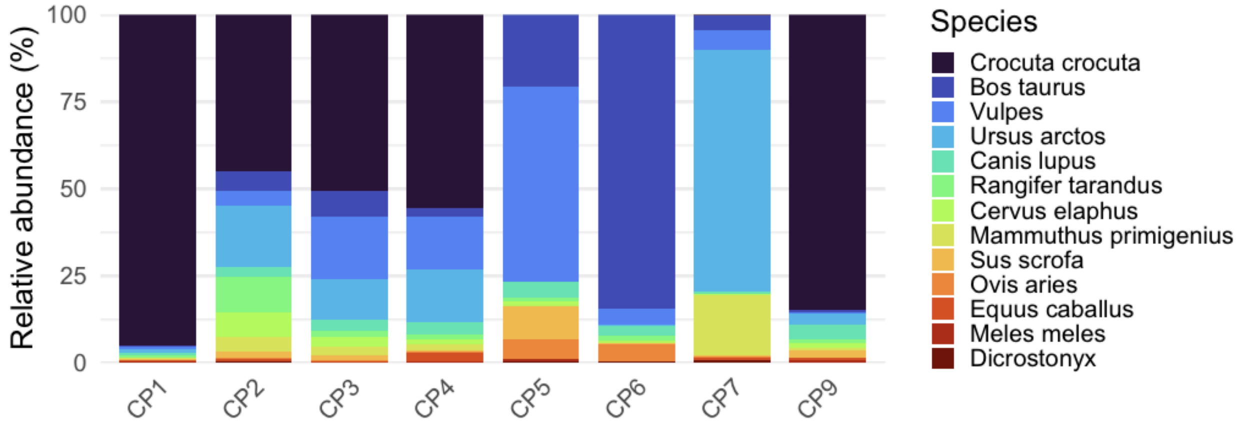


Supplementary Figure 17 - Taxonomic composition based on blast analysis of the full set of reads.

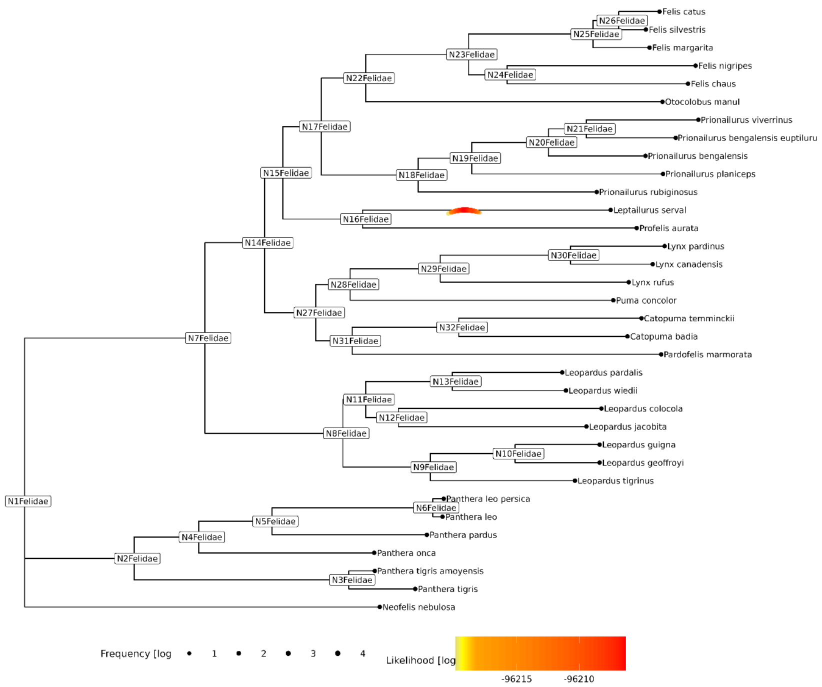


**Supplementary Figure 18** - Soibean assignment of sequences to *Leptailurus serval* (serval) within the Felidae mtDNA phylogenetic tree. We reassigned these reads to Crocuta crocuta using blast.

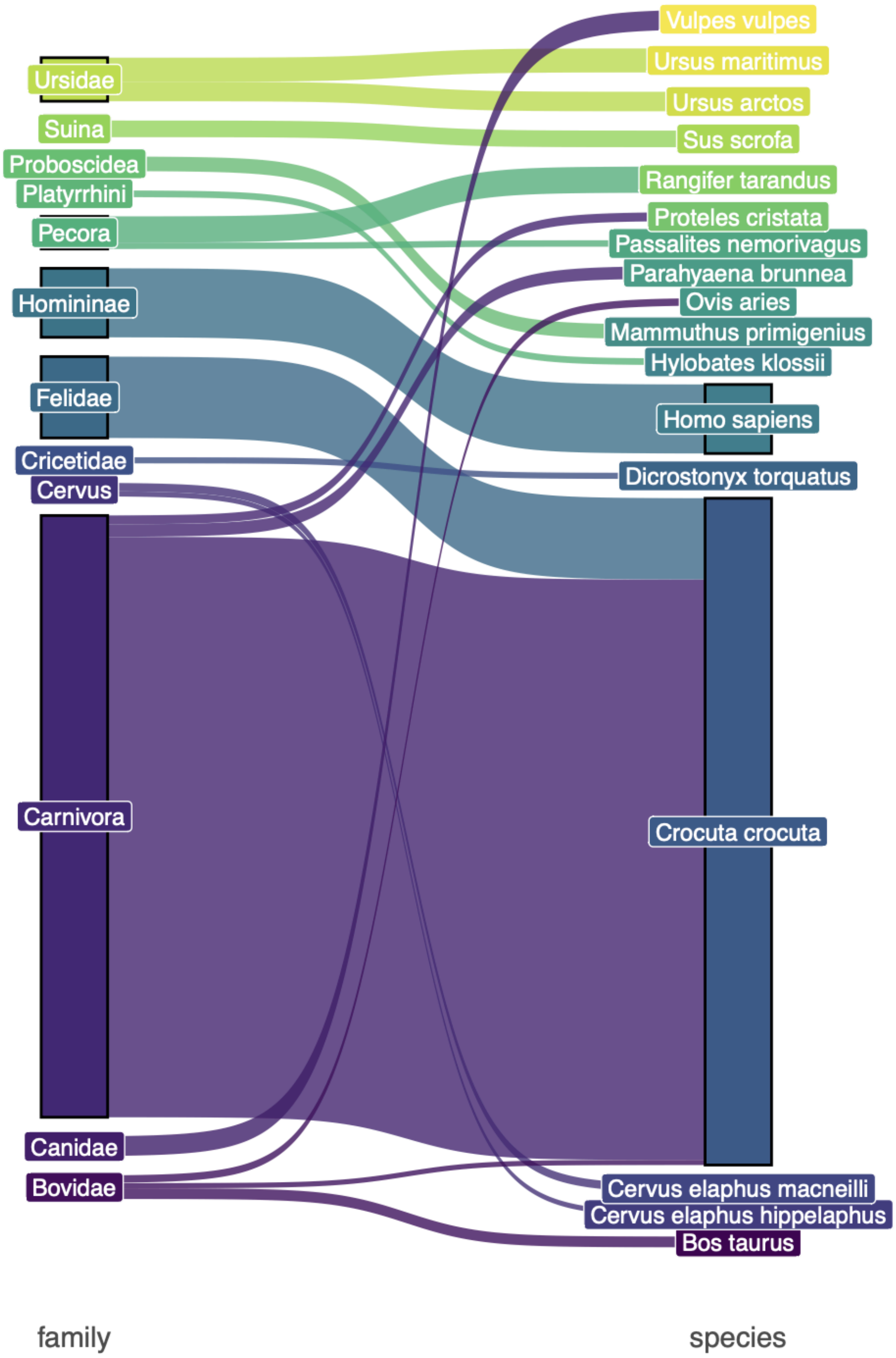


Supplementary Figure 19 - Comparison between euka family classification (left) and blast species classification (right). Line width is proportional to read count. Most of Felidae reads were shown to belong to Crocuta.

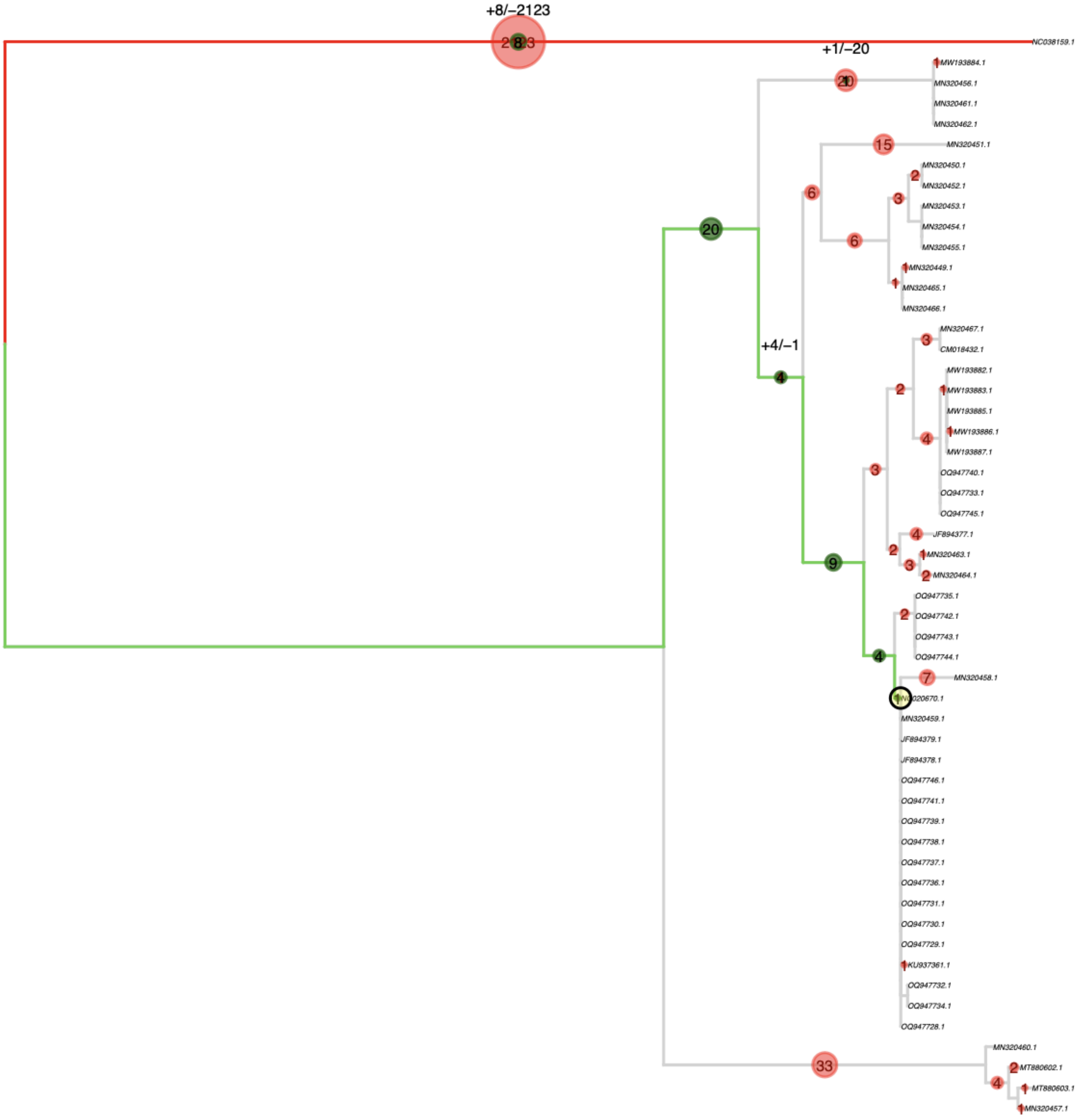


Supplementary Figure 20 - Phylogenetic placement of sequencing reads from HyL1 in a tree containing present-day and ancient hyena mitochondrial sequences. We obtained similar placements for samples from other layers.

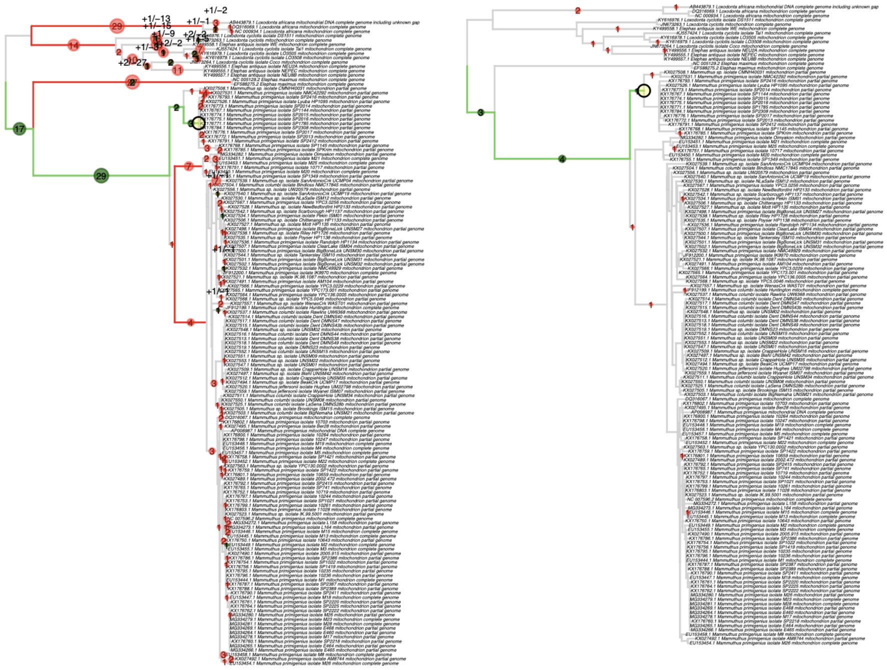

Supplementary Figure 21 - Phylogenetic placement of sequencing reads from HyL1 and EH2 in a tree containing ancient mammoth mitochondrial sequences. We obtained similar placements for samples from other layers.

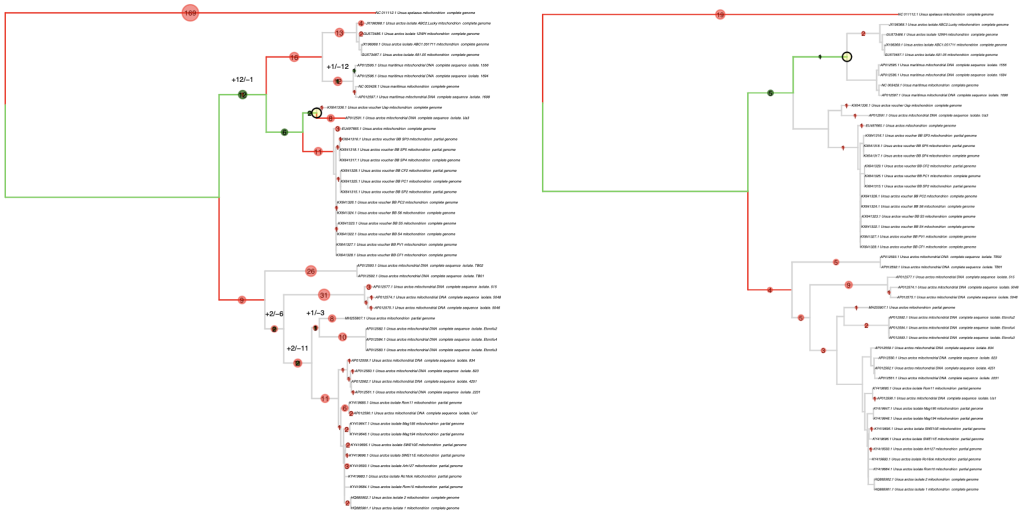

Supplementary Figure 22 - Phylogenetic placement of sequencing reads from the same layer (HyL2) in a tree containing bear mitochondrial sequences. Brown bear reads in HyL2 predominantly belong to clade 1b (A), but through competitive mapping we obtained a minority of reads from the same layer which belonged to clade 2 (B). Samples from other layers belonged to clade 1b.

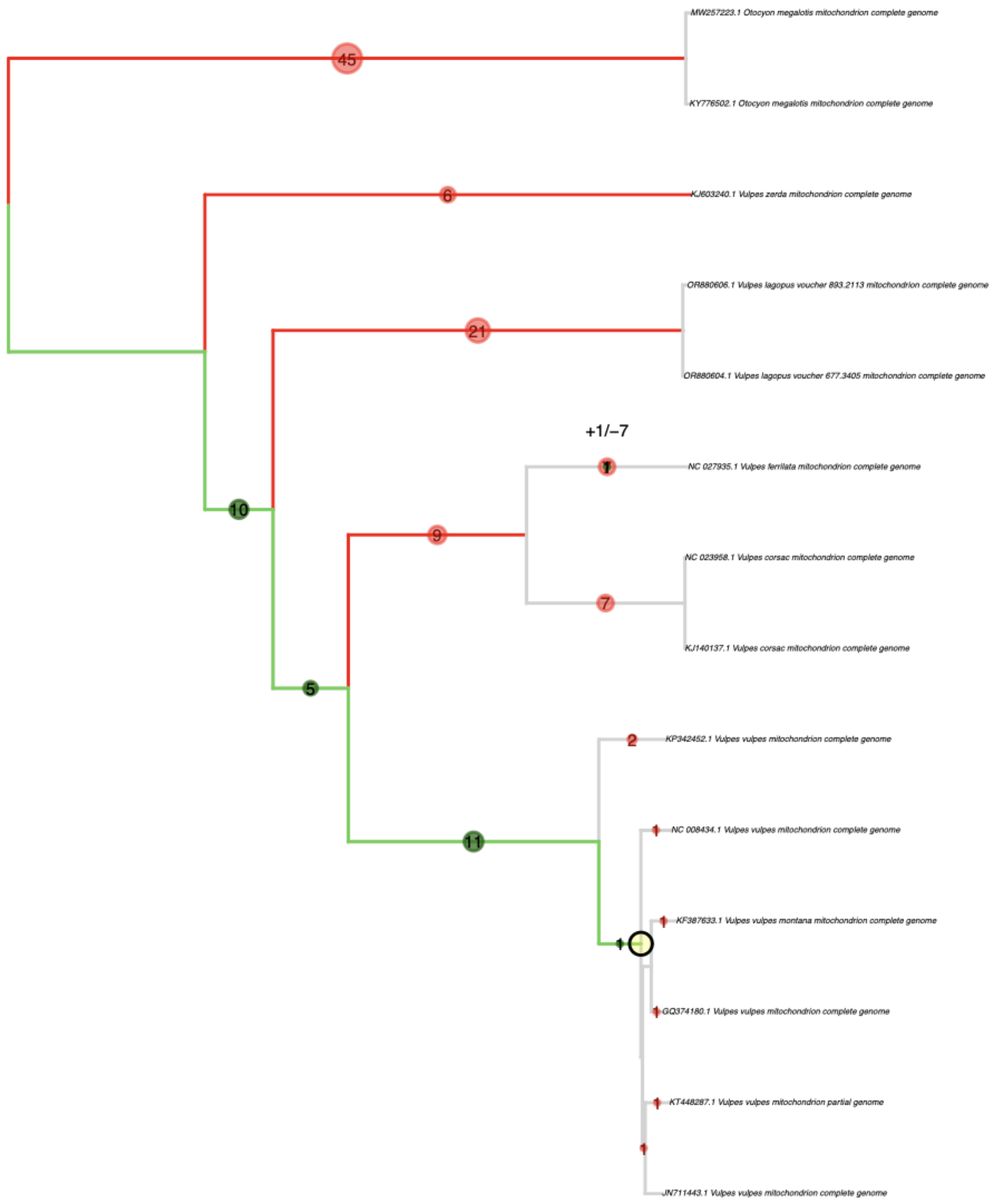


Supplementary Figure 23 - Phylogenetic placement of HyL3 demonstrating that the reads belong to red fox (*Vulpes vulpes*) and not Arctic fox (*Vulpes lagopus*).

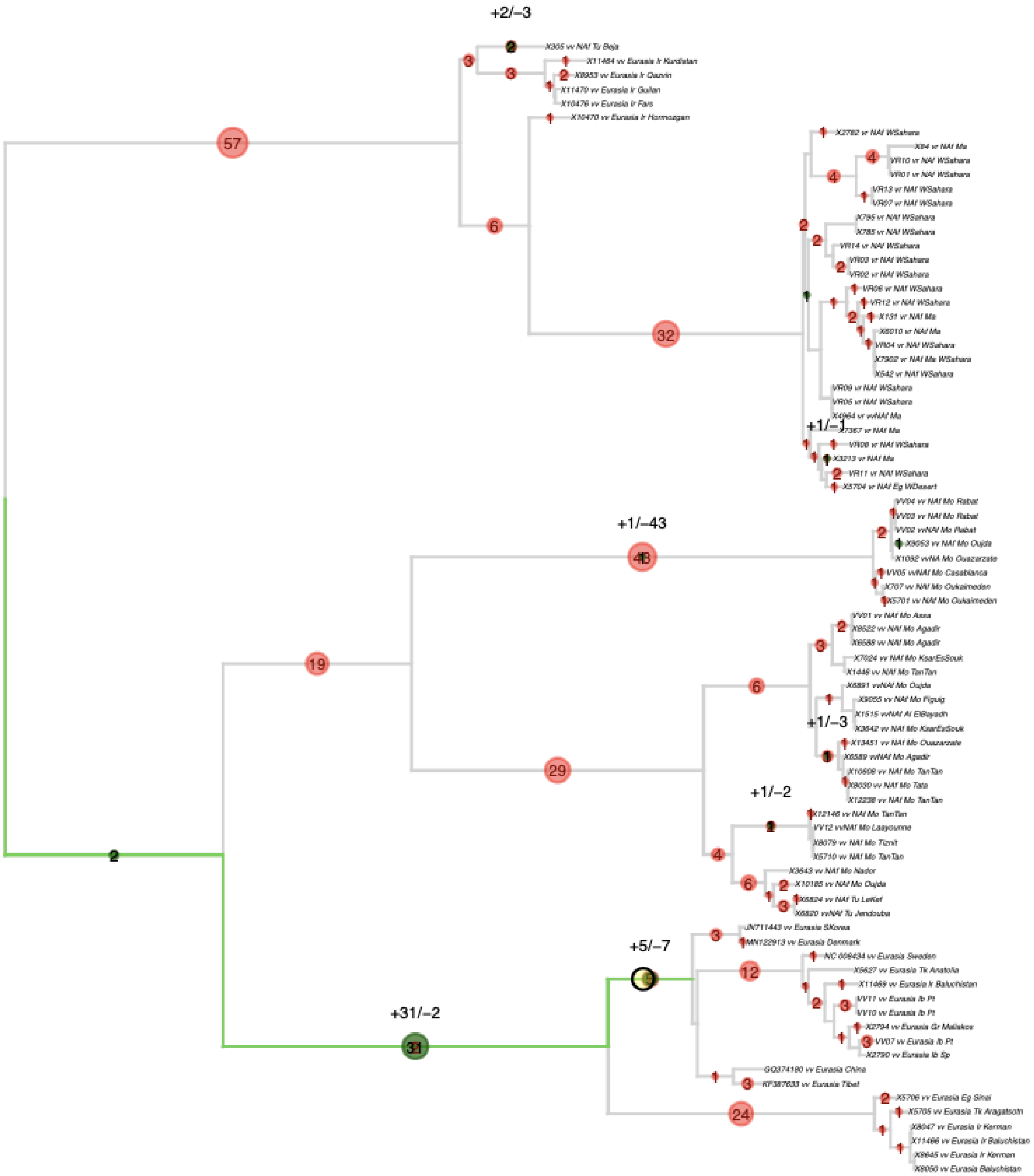


Supplementary Figure 24 - Phylogenetic placement of HyL3 in a tree built with present-day fox mitochondrial DNA, supporting membership to the Eurasian clade of red fox.

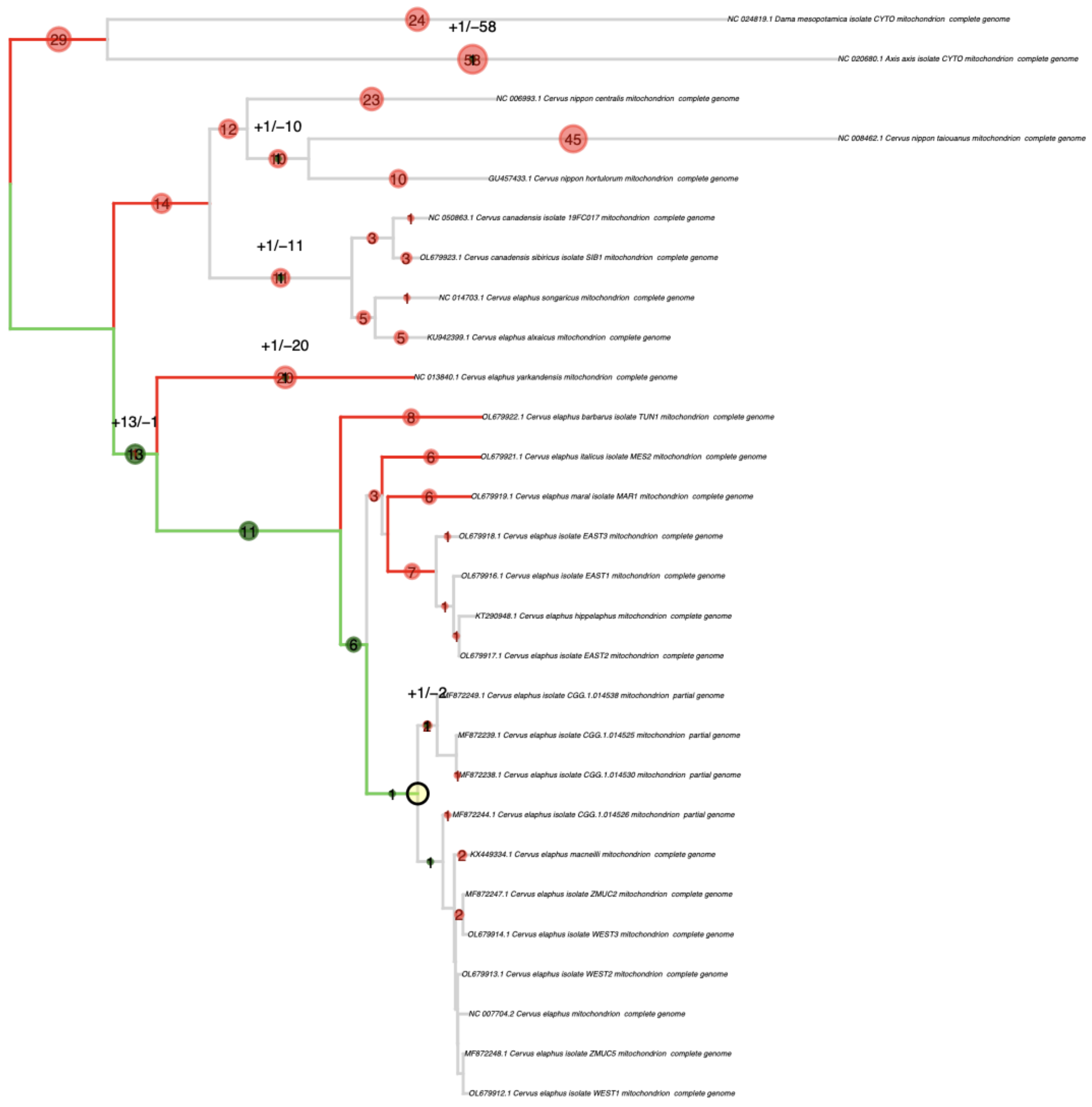


Supplementary Figure 25 - Phylogenetic placement of HyL2 in a tree built with present-day deer mitochondrial DNA, supporting membership to the Eurasian clade of red deer.

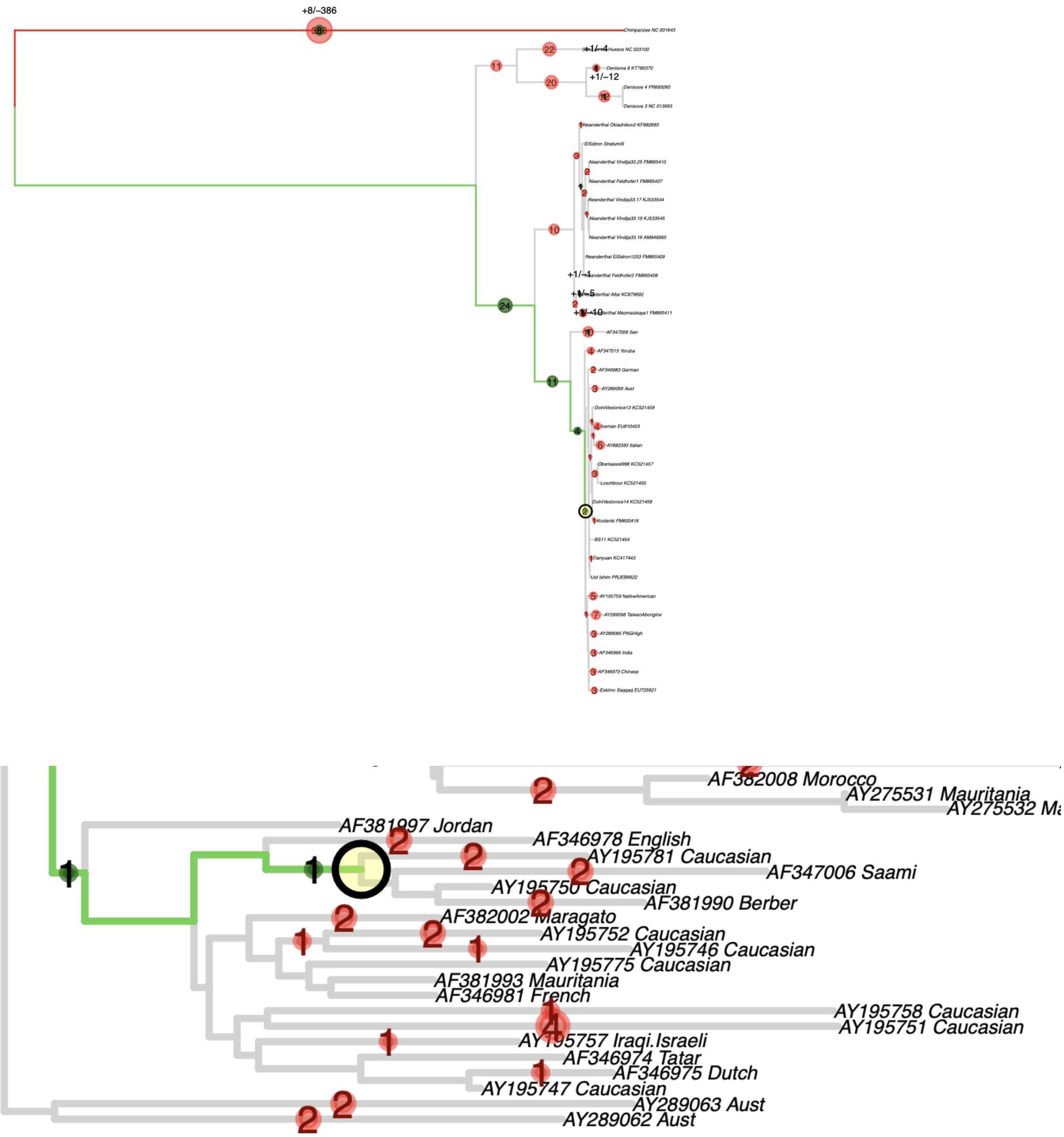


Supplementary Figure 26 - A) Phylogenetic placement of HyL1 in a tree composed of primate sequences, including chimpanzee, Neanderthals, Denisovans and modern humans, with clear membership in the latter. B) Placement of HyL in a modern human mitochondrial phylogeny, in the branch corresponding to the V haplogroup which includes samples AY195781 (hg V6b), AF347006 (V7a1c), AY195750 (V) and AF3811990 (V25).

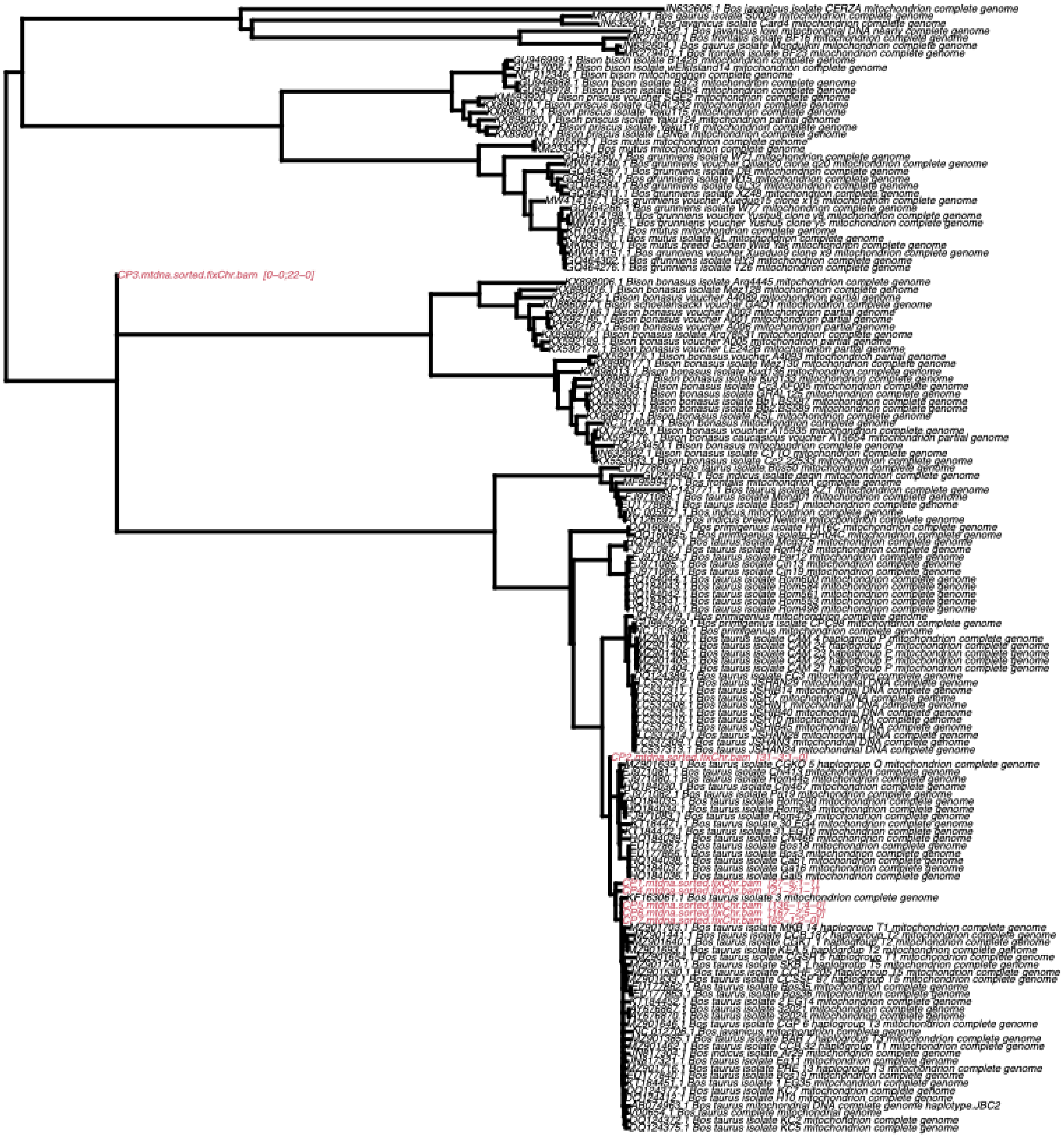

Supplementary Figure 27 - Phylogenetic placement of HyL2 in a tree built with present-day cattle mitochondrial DNA, supporting membership to the Eurasian clade of cattle.
